# The MHC requirement for αβ T-cell specification is heterochronically gated

**DOI:** 10.64898/2026.11.19.767775

**Authors:** Xin Sun, Feiyang Ma, Andrew J. Zak, Rachael Bogle, Luis O. Correa-Medero, Hatem G. Fallouh, Patrick T. Rudak, Syed M. Rizvi, Isabel O. Vanderzee, Crystal Schmerk, Li Chen, S. M. Mansour Haeryfar, Kathryn A. Eaton, Shannon A. Carty, Fei Wen, Allison C. Billi, Lam C. Tsoi, Johann E. Gudjonsson, Amale Laouar, Yasmina Laouar

## Abstract

Major histocompatibility complex (MHC)-dependent thymic selection is a defining principle of αβ T-cell development, yet mice lacking β2m-dependent MHC class I and MHC class II retain rare αβ T cells whose developmental significance has remained unresolved. Here we show that this residual compartment is not simply the product of failed selection. Single-cell and clonotypic profiling resolves distinct outcomes, including clonally restricted CD4-associated states and a diversified CD8⁺ αβ T-cell population that converges on physiological wild-type virtual memory. These CD8⁺ cells lack conventional self-MHC calibration, retain broad T-cell receptor (TCR) diversity, respond rapidly to T-cell receptor and cytokine stimulation, and confer protection during bacterial infection. Their preservation is developmentally restricted: the MHC-independent state carries a neonatal transcriptional and phenotypic imprint, is favored by fetal but not adult progenitors, and is reduced by neonatal Lin28 inhibition. While these data do not displace the central requirement for MHC in generating the conventional naive αβ T-cell repertoire, they establish that αβ T-cell specification can proceed during early life in its absence, yielding a physiological and functionally competent CD8⁺ αβ T-cell fate. Thus, the MHC requirement for αβ T-cell specification is heterochronically gated.

## INTRODUCTION

The association between αβ T-cell receptors and MHC molecules is a defining organizing principle of vertebrate adaptive immunity. Mature αβ T cells recognize antigen in the context of MHC or MHC-related molecules, a relationship shaped by both evolutionary constraints on TCR recognition and developmental selection in the thymus^1–4^. Thymic interactions between the TCR and self-peptide–MHC govern both selection and lineage fate, forming the basis of the naive peripheral αβ T-cell repertoire. MHC restriction is therefore both a property of mature αβ T cells and an outcome of the developmental process that produces them.

The developmental requirement for MHC was established unequivocally by genetic experiments. Disruption of β2-microglobulin (β2m), which is required for stable expression of MHC class I molecules, nearly abolishes mature CD8⁺ αβ T cells^5,6^, and disruption of MHC class II results in a comparable loss of mature CD4⁺ αβ T cells^7,8^. Combined deficiency of β2m- dependent MHC class I and MHC class II severely depletes both compartments^9^, but does not eliminate αβ T cells entirely, leaving a small residual peripheral population ^10–13^. These studies established MHC selection as the dominant pathway generating the naive CD4⁺ and CD8⁺ αβ T-cell compartments and became foundational evidence for the requirement of self-MHC recognition during thymic development^5–7,9^. That conclusion is correct, but incomplete. Those experiments were interpreted in the early 1990s when the peripheral αβ T-cell compartment was understood predominantly through its abundant naive CD4⁺ and CD8⁺ populations. In that framework, the decisive observation was numerical collapse: the major naive compartments failed to develop, whereas the much smaller population that remained was viewed against the magnitude of this loss. In more than three decades since, the landscape of αβ T-cell biology has changed substantially. Numerically minor populations are now recognized as coherent developmental and functional compartments, including invariant NKT cells, MAIT cells, regulatory T cells and virtual memory T cells^3,14–16^, arguing that rarity does not imply developmental insignificance. By the same reasoning, the residual αβ T cells in MHC-deficient mice may likewise represent biologically meaningful developmental outcomes whose significance has remained unrecognized.

Several observations make this possibility biologically plausible. αβ TCRs are now recognized to engage ligands beyond conventional peptide–MHC complexes^17–20^. Biochemical and structural studies of αβ TCRs from MHC- and coreceptor-deficient mice demonstrated the ability of αβ TCRs to directly engage folded native protein surfaces through their CDR-loop antigen-combining surface^19–21^. The capacity for direct protein recognition was also demonstrated in naturally occurring human and murine αβ TCRs. A human αβ TCR isolated from a renal-cell carcinoma–reactive T-cell line bound a protein complex lacking any MHC-related component and without requiring classical peptide processing^22^, and a murine αβ TCR directly bound intact phycoerythrin^23^. A second principle emerged from studies of T-cell development under impaired TCR signaling: disruption of the signaling machinery does not invariably prevent αβ T-cell development. Disruption of the Tec kinases ITK and RLK, or the ERK–ELK4/ELK1 signaling axis, demonstrated that αβ T-cell development can proceed toward an alternative innate-like fate when conventional thymic selection is impaired^24–27^.

Another key dimension in understanding the requirement for MHC is developmental time. Thymopoiesis unfolds within an epithelial environment that undergoes profound ontogenic remodeling, from an immature organization in early life to differentiated cortical and medullary compartments with established programs of antigen presentation and tissue-restricted antigen expression in adulthood^28^. Concurrently, hematopoietic progenitors undergo developmental transitions that reshape their lymphoid potential^29,30^, with early-life hematopoiesis preferentially generating T cells endowed with innate-like properties^31–35^. These coordinated changes in thymic environment and progenitor competence raise the possibility that the requirement for MHC in αβ T-cell development is itself developmentally regulated. The unresolved question is therefore not whether MHC is required to generate the dominant naive αβ T-cell repertoire—it is—but whether the αβ T cells that persist in its absence are stochastic remnants of failed conventional selection or the product of a distinct developmental program.

## RESULTS

### Aberrant and non-aberrant αβ-lineage fates

Residual αβ T cells have long been documented in MHC-deficient mice^9–13^, yet whether this compartment represents stochastic escape from failed selection or an organized set of alternative αβ-lineage states remains unresolved. To distinguish these possibilities, we mapped the residual αβ T-cell compartment in *β2m^⁻/⁻^MHCII^⁻/⁻^* mice at cellular, transcriptional and clonal resolution. Mass cytometry of the peripheral immune landscape visualized the contraction of the αβ T-cell compartment to a small residual population in *β2m^⁻/⁻^MHCII^⁻/⁻^* mice, and flow cytometry quantified this contraction, indicating that lack of MHC severely contracts but does not abolish αβ-lineage output (Fig. 1a,b; Extended Data Fig. 1a,b). No αβ T cells were detected in *Rag2^⁻/⁻^* or *Foxn1^⁻/⁻^* mice (Extended Data Fig. 1c), and neither *i*NKT nor MAIT cells were detected in *β2m^⁻/⁻^MHCII^⁻/⁻^* mice, providing stringent controls for the detection of the residual TCRβ⁺ population in *β2m^⁻/⁻^MHCII^⁻/⁻^* mice^3,16,36^ (Extended Data Fig. 1d,e).

**Fig. 1:**
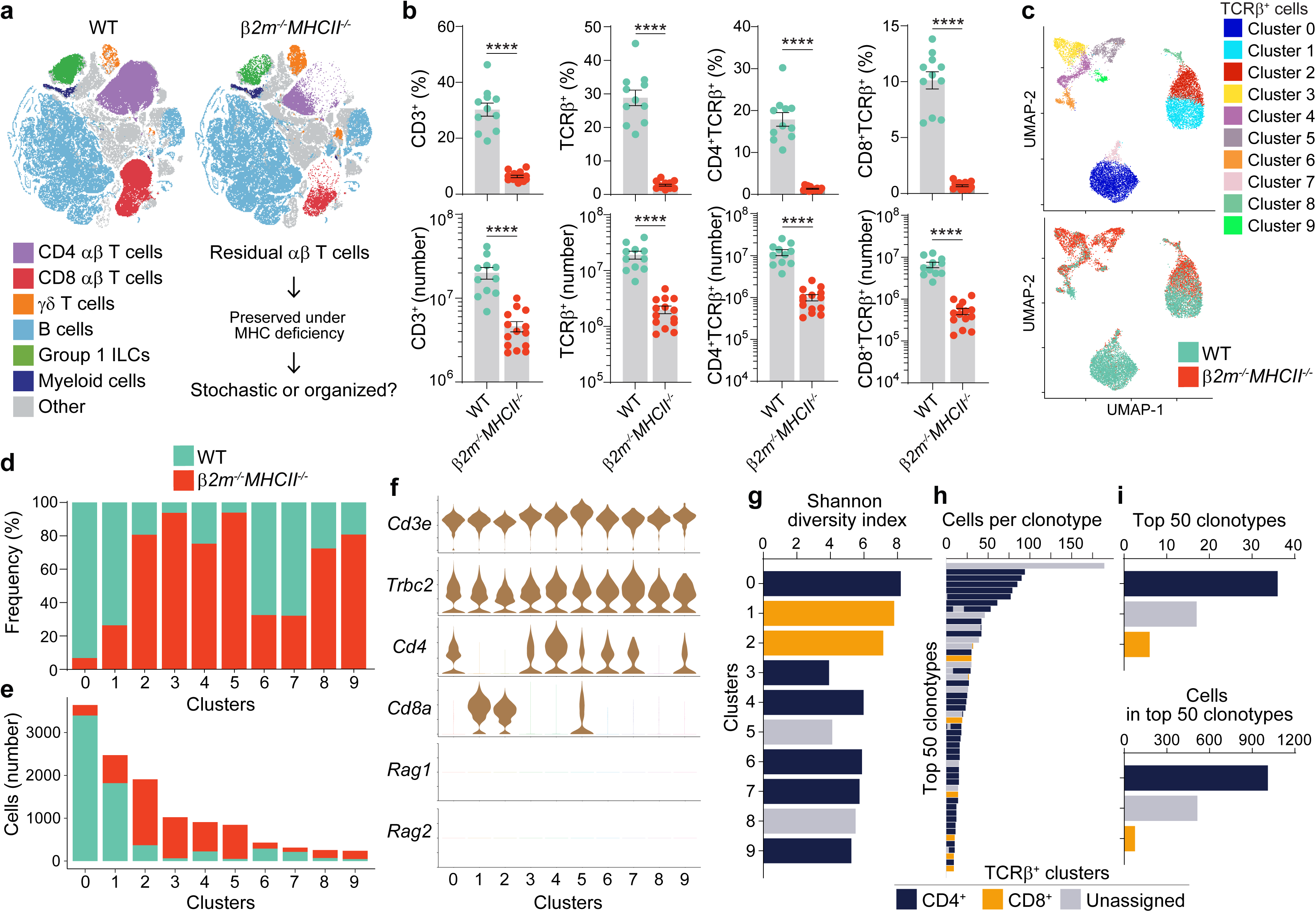
MHC deficiency reveals heterogeneity in residual αβ T-cell clonality. **a,** CyTOF analysis of WT and *β2m^⁻/⁻^MHCII^⁻/⁻^* splenocytes. t-SNE immune landscape annotated by major immune subset. **b,** Frequency and cell number of splenic CD3⁺, CD3⁺TCRβ⁺, CD4⁺TCRβ⁺ and CD8⁺TCRβ⁺ cells in WT and *β2m^-/-^MHCII^-/-^* mice. **c-i,** scRNA-seq and scTCR immune profiling analysis of splenic αβ T cells from WT and *β2m^⁻/⁻^MHCII^⁻/⁻^* mice. **c,** UMAP distribution of splenic αβ T cells by clusters (top) and genotypes (bottom). **d,** Genotype contribution to each cluster. **e,** Cell number of each genotype in each cluster. **f,** Violin plots showing *Cd3e*, *Trbc2*, *Cd4*, *Cd8a, Rag1,* and *Rag2* expression across clusters. **g,** Shannon diversity index of the TCR clone repertoire of each cluster. **h,** Cell number of 50 most enriched TCR clonotypes and the contribution from CD4 T cell clusters (Cluster 0, 3, 4, 6, 7, 9), CD8 T cell clusters (Cluster 1, 2), and unassigned clusters (Cluster 5, 8). **i,** Within 50 most enriched TCR clonotypes, the number of clonotypes (top) and the number of cells (bottom) grouped by CD4 T cell clusters, CD8 T cell clusters, and unassigned clusters. Data (**b**) are from 5-6 independent experiments; n = 11-14. Unpaired two-tailed t-test. Data are mean ± s.e.m.; \**p* < 0.05, \*\**p* < 0.01, \*\*\**p* < 0.001, \*\*\*\**p* < 0.0001; ns, not significant. See also Extended Data Fig. 1 and 2.

To resolve the identity of this residual compartment, we mapped it against the wild-type αβ T-cell landscape by integrating transcriptional identity with clonal architecture (Fig. 1c-i). Unsupervised clustering resolved ten clusters with a non-uniform distribution across genotypes, revealing that the residual αβ T-cell compartment in *β2m^⁻/⁻^MHCII^⁻/⁻^* mice is not a miniature wild- type landscape. Wild-type CD4^+^ and CD8^+^ αβ T cells were predominantly represented by clusters 0 and 1, respectively. Against this reference, *β2m^⁻/⁻^MHCII^⁻/⁻^* CD8^+^ αβ T cells converged on cluster 2, whereas the remaining αβ T cells dispersed across multiple CD4 and unassigned states that were either scarce or virtually absent from the wild-type landscape (Fig. 1d-f). Paired TCR profiling further showed that the residual αβ T-cell compartment in *β2m^⁻/⁻^MHCII^⁻/⁻^* mice was not clonally uniform. CD4 and unassigned states persisting in *β2m^⁻/⁻^MHCII^⁻/⁻^* mice exhibited a lower Shannon diversity index and dominated the pool of the 50 most abundant clonotypes. In contrast, the persisting CD8 state retained clonotype diversity comparable to wild-type CD8⁺ cells and was minimally represented among the expanded clonotypes, arguing against a model in which loss of MHC inevitably constrains αβ-lineage output to clonally restricted residual states (Fig. 1g-i).

To rule out the possibility that CD8⁺ αβ T cells from *β2m^⁻/⁻^MHCII⁻^/⁻^* mice were calibrated by trace MHC signals during development, we used mixed lymphocyte reactions as a functional readout of thymic MHC calibration (Extended Data Fig. 2). In conventionally selected CD8^+^ αβ T cells, thymic exposure to self MHC establishes tolerance to syngeneic MHC while preserving responsiveness to allogeneic MHC^10,37^. In contrast, CD8^+^ αβ T cells from *β2m^⁻/⁻^MHCII^⁻/⁻^* mice proliferated in response to both B6 background-matched and BALB/c allogeneic MHC-expressing stimulators, indicating that they were not calibrated by self MHC in the manner that restrains syngeneic MHC reactivity in conventionally selected CD8⁺ αβ T cells. Thus, exposure to trace MHC, if any, was insufficient to impose conventional MHC calibration on the CD8⁺ αβ T cells persisting in *β2m^⁻/⁻^MHCII^⁻/⁻^* mice.

### A physiological non-naive fate

Persistence of a CD8⁺ αβ T-cell state with preserved TCR clonotype diversity despite the absence of MHC calibration raised a fundamental question: whether this population represents an aberrant output or a physiological CD8 program that remains accessible without MHC instruction. Relative to the naive program, cluster 2 expressed *Tbx21*, *Eomes*, *Bcl2*, *Il2rb*, *Cxcr3* and *Cd44*, together with low *Itga4*, defining the canonical molecular axis of virtual memory^14,38^ (Fig. 2a-c). This identity was coupled to a pre-armed effector architecture. Cytotoxic genes *Prf1*, *Gzmb* and *Serpinb9*, effector regulators *Runx3* and *Zeb2*, and interferon-associated genes *Ifng*, *Ifngr1* and *Mx1* were enriched. Genes supporting cytotoxic execution were likewise enriched across immunological-synapse organization (*Adap1*, *Anxa2*, *Pld2*)^39,40^, vesicular trafficking and cytoskeletal remodeling (*Rhoc*, *Myo1f*, *S100a4*, *Lyst)* and Ca²⁺ signaling (*Trpc1*, *Atp2b4*)^41^. Enrichment of *Il12rb1*, *Il12rb2*, *Il18r1* and *Il15ra*, together with *Klrk1*, *Klrc1*, *Klrc2* and multiple *Klra* family genes, further defined an innate-responsive module. Pathway analysis independently placed cluster 2 at the intersection of CD8⁺ T-cell memory and NK-cell cytotoxicity, clearly distinguishing this state from the quiescent naive program (Fig. 2d).

**Fig. 2:**
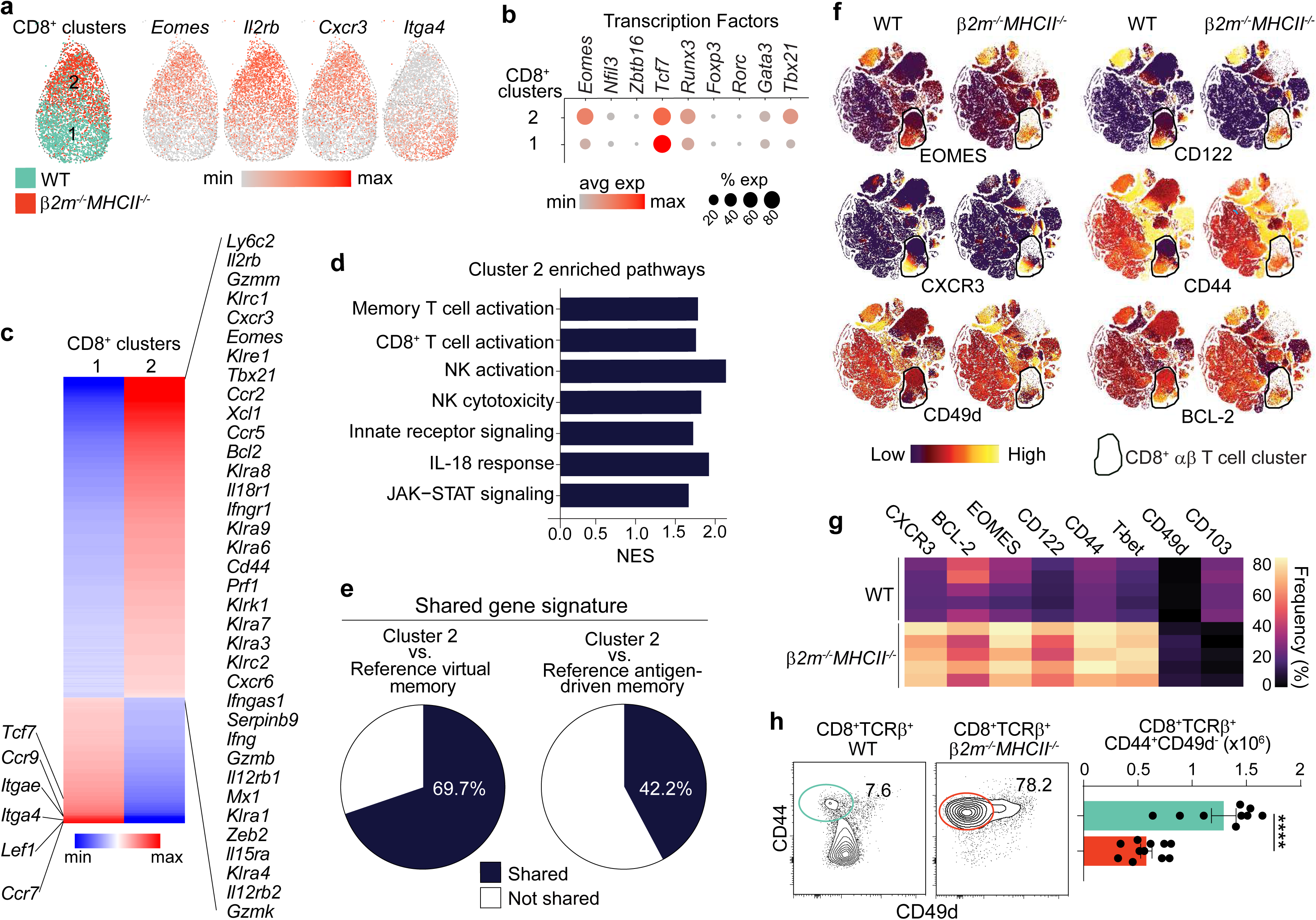
CD8⁺ αβ T cells from *β2m^⁻/⁻^MHCII^⁻/⁻^* mice converge on physiological virtual memory. **a-e,** scRNA-seq analysis of splenic αβ T cells from WT and *β2m^⁻/⁻^MHCII^⁻/⁻^* mice in Fig. 1c. **a,** UMAP distribution of CD8⁺ αβ T-cell clusters by genotype, with clusters 1 and 2 delineated (left). Feature plots of indicated genes in clusters 1 and 2 (right). **b,** Dot plot of indicated transcription factor gene expression in clusters 1 and 2. **c,** Heatmap of the 778 DEGs identified between clusters 1 and 2; selected genes are labeled. **d,** Selected GSEA pathways enriched in cluster 2 relative to cluster 1. NES, normalized enrichment score. **e,** Cross-dataset validation of cluster 2 versus cluster 1 DEGs against reference virtual memory T cells and reference antigen-driven memory versus naive CD8⁺ T-cell DEGs (GSE112304; virtual memory T cells, CD3⁺CD8⁺CD44⁺CD49d⁻; antigen-driven memory, CD3⁺CD8⁺CD44⁺CD49d⁺; naive, CD3⁺CD8⁺CD44⁻)^42^. Pie chart shows overlap of 293/420 reference virtual memory T cells DEGs and 336/796 antigen-driven memory T cells DEGs with the cluster 2 DEG set. **f,g,** CyTOF analysis of WT and *β2m^⁻/⁻^MHCII^⁻/⁻^* splenocytes in Fig 1a. **f,** Relative expression of indicated markers across the immune landscape; black gate delineates the CD8⁺ αβ T-cell cluster. **g,** Heatmap of indicated markers frequencies within the CD8^+^ αβ T-cell cluster from WT and *β2m^-/-^MHCII^-/-^* splenocytes; n = 5 per genotype. **h,** Flow cytometry analysis of WT and *β2m^⁻/⁻^MHCII^⁻/⁻^* splenocytes. CD44/CD49d distribution among CD8⁺ αβ T cells (left), with the cell number of CD44⁺CD49d⁻ CD8 T cells (right). Data (**h**) are from 4-6 independent experiments; n = 9-11. Unpaired two-tailed t-test. Data are mean ± s.e.m.; \**p* < 0.05, \*\**p* < 0.01, \*\*\**p* < 0.001, \*\*\*\**p* < 0.0001; ns, not significant. See also Extended Data Fig. 3-9.

Cross-referencing the cluster 2 signature against public-dataset profiles^42^ revealed 69.7% overlap with wild-type virtual memory, compared with 42.2% overlap with antigen-driven memory, anchoring the residual *β2m^⁻/⁻^MHCII^⁻/⁻^* population to a physiological virtual memory state (Fig. 2e). The program not shared with antigen-driven memory includes lineage and survival modules (*Runx3*, *Id2*, *Bcl2*, *Il15ra*), reduced chromatin and anabolic remodeling (*Tet1*, *Tet3*, *Chd2*, *Srebf2*, *Dhcr24*), and a distinct type I interferon-response program (reduced *Stat1* with increased *Ifit3*, *Oas3* and *Usp18*), distinguishing CD8⁺ αβ T cells arising without MHC calibration from the adaptive remodeling associated with antigen-driven differentiation^43–46^ (Extended Data Fig. 3). Because γδ T cells are preserved in *β2m^⁻/⁻^MHCII^⁻/⁻^* mice and can provide a source of IL-4^47–51^, we investigated whether the residual CD8⁺ αβ T cells correspond to an IL-4-induced memory-like state^52^. We therefore induced a memory-like CD8⁺ αβ T-cell population through IL-4 administration *in vivo* and compared its transcriptional profile with that of *β2m^⁻/⁻^MHCII^⁻/⁻^* CD8⁺ αβ T cells. Despite a shared core virtual memory module, the two populations diverged extensively across ∼2,700 genes, with *β2m^⁻/⁻^MHCII^⁻/⁻^* cells lacking the prominent inducible activation signature of IL-4-induced memory-like cells (Extended Data Fig. 4).

Protein-level mapping reproduced this identity. CITRUS resolved the CD8⁺ αβ T-cell nodes into a wild-type naive branch, with negligible *β2m^⁻/⁻^MHCII^⁻/⁻^* representation, and a virtual memory branch (EOMES⁺BCL-2⁺CD122⁺CXCR3⁺CD44⁺CD49d^lo^) shared between wild-type and *β2m^⁻/⁻^MHCII^⁻/⁻^* cells (Extended Data Fig. 5). t-SNE showed that this preserved state occupies only a small region of the wild-type CD8⁺ αβ T-cell landscape, most of which corresponds to the naive state depleted in *β2m^⁻/⁻^MHCII^⁻/⁻^* mice (Fig. 2f,g). Flow cytometry reproduced this virtual memory phenotype across ∼30 markers and quantified a residual pool of ∼0.5 × 10⁶ cells in *β2m^⁻/⁻^MHCII^⁻/⁻^* mice that remained numerically stable over a 1-year period (Fig. 2h; Extended Data Fig. 6, 7).

We next asked whether the correspondence between wild-type and *β2m^⁻/⁻^MHCII^-/-^* virtual memory cells extended to their TCR repertoire architecture. Within the transcriptionally defined virtual memory state (cluster 2), both populations exhibited broad and largely concordant TRAV, TRAJ, TRBV, TRBD and TRBJ usage, sharing the dominant gene segments represented in the physiological wild-type repertoire (Extended Data Fig. 8a). These similarities were also evident in the CDR3 regions of both α- and β-chains, which displayed closely matched amino-acid composition and positional entropy (Extended Data Fig. 8b,c). Shannon diversity analysis further revealed substantial repertoire diversity in *β2m^⁻/⁻^MHCII^-/-^* virtual memory cells, comparable to their wild-type counterparts (Extended Data Fig. 8d). Bulk TCRβ sequencing and Vβ profiling by flow cytometry corroborated the shared repertoire architecture of wild-type and *β2m^⁻/⁻^MHCII^-/-^* virtual memory cells (Extended Data Fig. 9). Thus, the preserved CD8⁺ αβ T-cell state not only shares the molecular identity of physiological virtual memory but also retains a broadly diversified receptor architecture closely resembling that of its wild-type counterpart. Because memory-like CD8⁺ αβ T-cell outputs can arise from TCR-signaling perturbation^24–27,53^, we tested whether CD8⁺ αβ T cells arising in the absence of MHC preserve the transcriptional architecture of this pathway or instead reflect disruption. Signaling components whose disruption is known to bias CD8⁺ αβ T- cell development toward memory-like states—including *Itk*, *Txk*, *Lcp2*, *Elk1* and *Elk4*—were preserved, with no differential enrichment of TCR-signaling gene programs between wild-type and *β2m^⁻/⁻^MHCII^⁻/⁻^* virtual memory cells, arguing against broad TCR-signaling disruption as the basis for preservation of this state under MHC deficiency (Extended Data Fig. 8e,f). Thus, CD8⁺ αβ T cells preserved in the absence of MHC are neither naive nor antigen-driven or IL-4-induced memory-like cells; they are a diversified, coherent population that converges transcriptionally, phenotypically and clonally on a physiological virtual memory state.

### Effector competence without priming

A defining prediction of this identity is rapid effector mobilization. TCR ligation elicited dose- dependent IFNγ production, with no measurable gain from CD28 co-stimulation, indicating that TCR engagement alone was sufficient to mobilize the rapid effector response associated with immunological memory (Fig. 3a). Bypassing the TCR with PMA–ionomycin induced robust IFNγ within 4 h, whereas IL-12 plus IL-18 elicited strong IFNγ production within 24 h (Fig. 3b). This pre- armed state extended to cytotoxic effector machinery. IL-15 induced granzyme B within 24 h *ex vivo*, and systemic poly(I:C) mobilized granzyme B *in vivo* within 18 h (Fig. 3c). In wild-type mice, the poly(I:C)-responsive granzyme B⁺ population mapped to the minor pre-existing virtual memory compartment, providing a physiological counterpart to the residual *β2m^⁻/⁻^MHCII^⁻/⁻^* CD8⁺ αβ T-cell pool (Extended Data Fig. 10a). Against LCMV-specific memory, *β2m^⁻/⁻^MHCII^⁻/⁻^* CD8⁺ αβ T cells retained the CD122^hi^BCL-2^hi^EOMES^hi^ profile characteristic of wild-type virtual memory rather than antigen-driven memory (Extended Data Fig. 10b). A critical question was whether this pre-armed effector state could provide meaningful host protection. Within 48 h of *Listeria monocytogenes* infection, *β2m^⁻/⁻^MHCII^⁻/⁻^* CD8⁺ αβ T cells mounted IFNγ responses comparable to wild-type virtual memory cells (Fig. 3d). Transfer into *Rag2^⁻/⁻^Il2rγ^⁻/⁻^* recipient mice established their protective capacity, reducing bacterial burdens in spleen and liver within 72 h to levels comparable to those achieved by wild-type virtual memory cells (Fig. 3e). Thus, MHC loss does not uniformly compromise CD8⁺ αβ T-cell biology; it selectively preserves a state capable of early host defense.

**Fig. 3:**
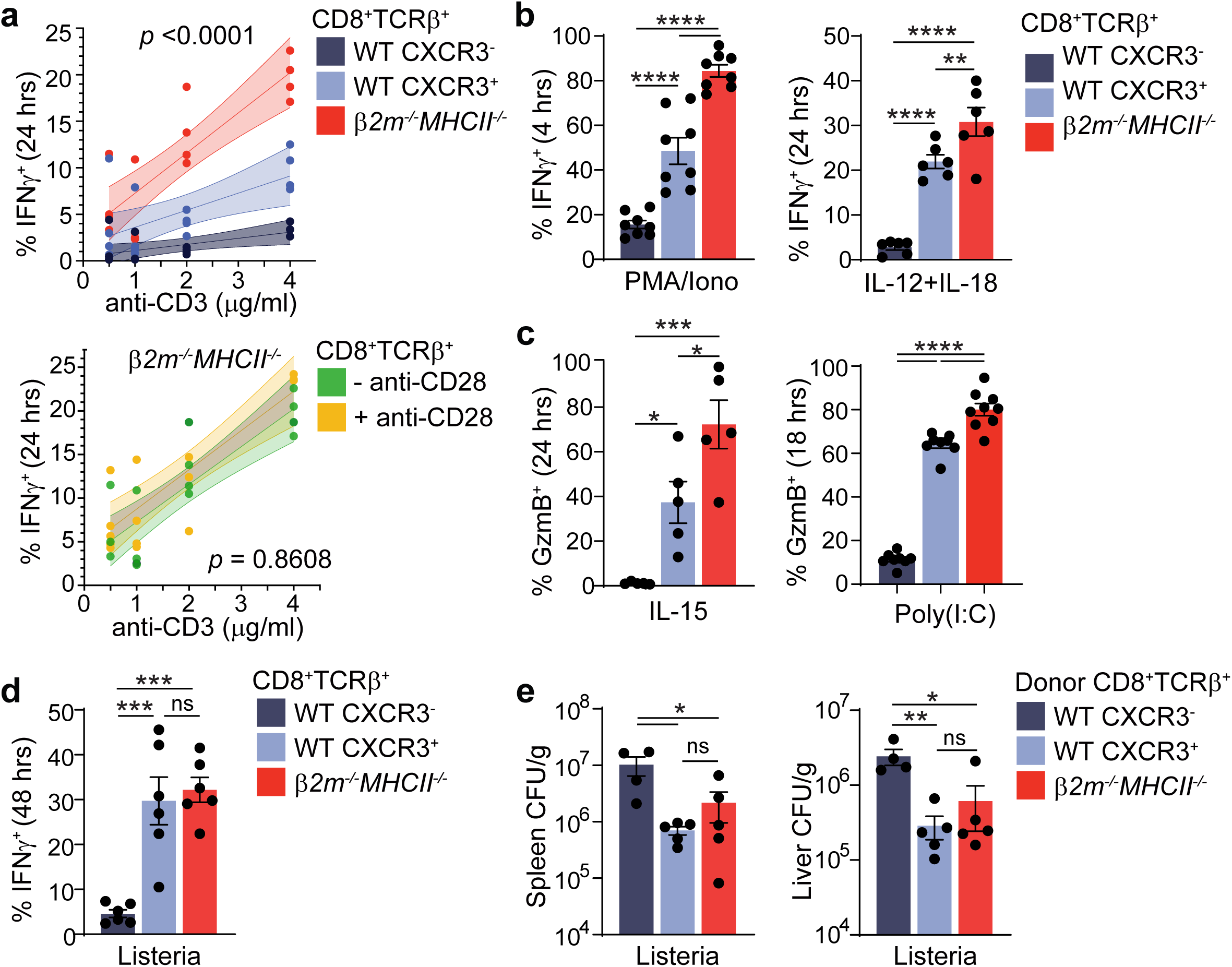
CD8⁺ αβ T cells from *β2m^⁻/⁻^MHCII^⁻/⁻^* mice are poised for immediate host defense. Function analysis among splenic WT CXCR3^-^ CD8 T cells, WT CXCR3^+^ CD8 T cells, and *β2m^⁻/⁻^MHCII^⁻/⁻^* CXCR3^+^ CD8 T cells. **a,** IFNγ⁺ frequency among indicated CD8⁺ αβ T-cell subsets after plate-bound anti-CD3 stimulation for 24 h (top), with or without soluble anti-CD28 co-stimulation (1 μg/ml) (bottom) ex vivo. **b,** IFNγ⁺ frequency among indicated CD8⁺ αβ T-cell subsets after PMA (50 ng/ml) plus ionomycin (1 μg/ml) stimulation for 4 h (left) or IL-12 (5 ng/ml) plus IL- 18 (25 ng/ml) stimulation for 24 h (right) ex vivo. **c,** GzmB^+^ frequency among indicated CD8⁺ αβ T-cell subsets after 24 h stimulation with IL-15 (50 ng/ml) ex vivo (left) or 18 h after *in vivo* poly(I:C) challenge (right). **d,** IFNγ⁺ frequency among indicated CD8⁺ αβ T-cell subsets 48 h after i.p. infection with 10^5^ CFU *L. monocytogenes*. **e,** Bacterial burden 72 h after *i.v.* infection with 10^4^ CFU *L. monocytogenes* in *Rag2^⁻/⁻^Il2rγ^⁻/⁻^* recipients transferred with the indicated CD8⁺ αβ T-cell subsets. Data are from 2–5 independent experiments; n = 4-9. For panel a, lines indicate simple linear regression with 95% confidence bands. P values indicate ANCOVA comparisons of slopes between groups. For (**b-e**), one-way ANOVA with Holm-Šídák’s test. Data are mean ± s.e.m.; \**p* < 0.05, \*\**p* < 0.01, \*\*\**p* < 0.001, \*\*\*\**p* < 0.0001; ns, not significant. See also Extended Data Fig. 10.

### Heterochronic specification

In *β2m^⁻/⁻^MHCII^⁻/⁻^* mice, CD8⁺ αβ T-cell development was not completely abolished, nor did it collapse into stochastic remnants, but instead resolved into a polyclonal, pre-armed virtual memory state, raising a broader biological question: under what physiological circumstances would such a fate make biological sense? Early ontogeny provided such a context. Analysis of a developmental single-cell atlas of non-hematopoietic thymic cells revealed that fetal and neonatal stroma expressed markedly lower levels of the classical MHC-I genes *B2m*, *H2-K1* and *H2-D1* and the MHC-II genes *H2-Aa*, *H2-Ab1* and *H2-Eb1* than adult stroma^28^. Central-tolerance machinery was similarly developmentally staged, with Aire^54^, Fezf2^55^ and multiple tissue-restricted antigens sparse during early ontogeny, consistent with a physiological period in which the molecular architecture supporting MHC calibration might be quantitatively attenuated (Extended Data Fig. 11, 12). CD8⁺ αβ T-cell output changes in parallel. Lineage tracing established that early- life thymopoiesis is strongly biased toward virtual memory output^56^, a fate promoted by the Lin28b–let-7 heterochronic axis through Eomes-dependent innate-like specification^57–60^. Whether these two developmental transitions converge to alter the MHC requirements of CD8⁺ αβ T-cell specification was unknown. Such a model predicts that MHC loss should not abolish all αβ T-cell specification but selectively preserve the early-life component of virtual memory. We tested this model across orthogonal developmental readouts, from single-cell ontogeny to precursor competence and heterochronic control (Fig. 4). Single-cell profiling first asked which ontogenic layer of virtual memory is preserved in the absence of MHC. The reference naive population was concentrated in C1 and C3, whereas virtual memory cells resolved into three additional states, C0, C2 and C4. When *β2m^⁻/⁻^MHCII^⁻/⁻^* CD8⁺ αβ T cells were benchmarked against these reference states, they mapped predominantly to C0 and C4 while largely excluding C2, indicating that MHC deficiency does not preserve the full spectrum of the virtual memory fate (Fig. 4a,b and Extended Data Fig. 13). What distinguishes the virtual memory states preserved in the absence of MHC from those that are lost? To determine whether that distinction tracks with developmental origin, we projected the retained and lost virtual memory states onto a neonatal transcriptional reference defined by neonatal versus adult-timestamped CD8⁺ αβ T cells^56^. C0 and C4 were strongly enriched for the neonatal program, whereas C2 showed substantially weaker neonatal imprint, placing the states preserved in *β2m^⁻/⁻^MHCII^⁻/⁻^* mice at the early-life end of the ontogeny axis (Fig. 4c). A neonatal gene score further quantified this hierarchy, placing wild-type virtual memory at an intermediate position consistent with a mixed developmental origin arising from continuous thymic output whereas CD8⁺ αβ T cells from adult *β2m^⁻/⁻^MHCII^⁻/⁻^* mice exhibited the highest neonatal score, consistent with an early-life origin (Fig. 4d,e). We demonstrated this developmental bias using FTOC and RTOC organ cultures, showing that EOMES⁺ CD8⁺ thymic output without MHC is favored with fetal progenitors but lost with adult progenitors (Fig. 4f). The associated pre-armed phenotype was already evident in neonatal *β2m^⁻/⁻^MHCII^⁻/⁻^* CD8⁺ αβ T cells and remained prominent in adulthood, as reflected by comparable granzyme B expression (Fig. 4g). We next tested whether this temporal restriction is heterochronically controlled. Neonatal Lin28 inhibition in *β2m^⁻/⁻^MHCII^⁻/⁻^* mice reduced Lin28b protein expression in T-cell precursors and contracted the virtual memory output, functionally linking this MHC-independent CD8⁺ αβ T-cell fate to the early-life developmental clock (Fig. 4h,i).

**Fig. 4:**
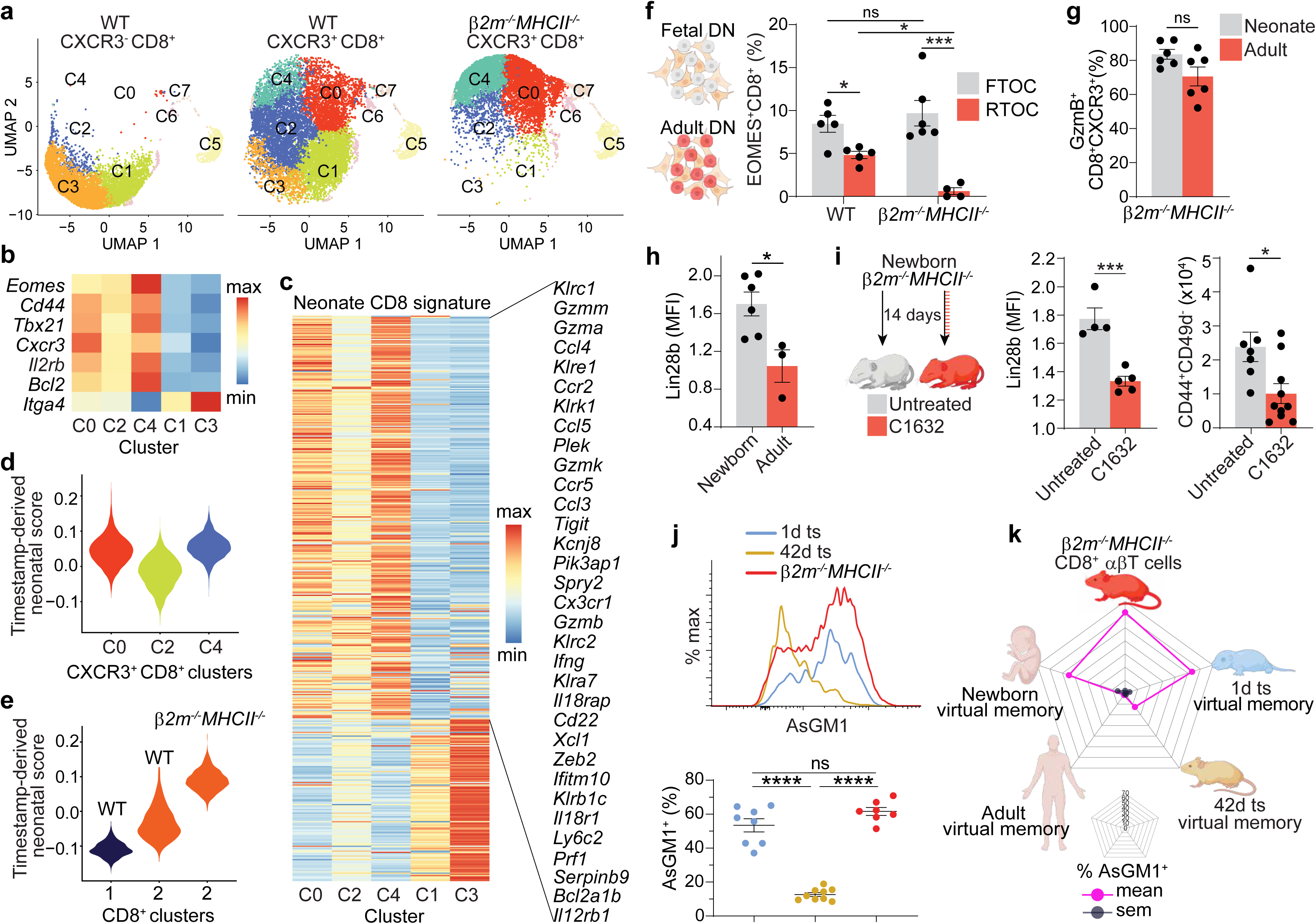
Early-life ontogeny gates MHC-independent CD8⁺ αβ T-cell specification. **a,b,** scRNA-seq analysis of splenic WT CXCR3⁻CD8⁺, WT CXCR3⁺CD8⁺, and *β2m^⁻/⁻^MHCII^⁻/⁻^* CXCR3⁺CD8⁺ αβ T cells. **a,** UMAP distribution by sorted CD8⁺ αβ T-cell population. **b,** Virtual memory-associated gene expression across clusters. **c-e,** Projection of a neonatal CD8 signature reference onto scRNA-seq datasets. The reference was defined from neonate versus adult timestamped CD8⁺ αβ T cells in GSE97795^56^. **c,** Heatmap of the 462 neonatal CD8 signature DEGs across the indicated clusters; selected genes are labeled. **d,e,** Timestamp-derived neonatal score, calculated from the 330 neonatal-enriched genes, across clusters from the dataset shown in Fig 4a (**d**), and across cluster 1 and cluster 2 from Fig 2a (**e**). **f,** EOMES⁺ frequency among CD8 single-positive thymocytes generated from fetal or adult DN thymocytes after 8 days of FTOC or RTOC. **g,** GzmB⁺ frequency among CD8⁺CXCR3⁺ αβ T cells from neonatal and adult *β2m^⁻/⁻^MHCII^⁻/⁻^* mice 18 h after *in vivo* poly(I:C) challenge. **h,** Lin28b protein expression in DN thymocytes from newborn and adult *β2m^⁻/⁻^MHCII^⁻/⁻^* mice, normalized to adult WT DN thymocytes. **i,** *β2m^⁻/⁻^MHCII^⁻/⁻^* dams were untreated or treated daily with the Lin28b inhibitor C1632 for 2 weeks postnatally. Lin28b protein expression in DN thymocytes at day 7 and splenic CD44^+^CD49d^-^ CD8 T cell number at day 14 were compared in neonates from untreated and C1632-treated dams. **j,** AsGM1 expression histogram overlay of indicated virtual memory T cell populations (top). AsGM1⁺ frequency among indicated virtual memory T cell populations (bottom). **k,** Radar plot of AsGM1⁺ frequency among virtual memory T cells from the indicated mouse and human sources. Pink lines indicate mean frequency and grey lines indicate s.e.m.; concentric pentagons indicate 10% frequency increments. Data (**f-k**) are from 2-6 independent experiments; n = 3-10. Two-way ANOVA with Newman–Keuls test (**f**); unpaired two-tailed t-test (**g-i**); One-way ANOVA with Tukey’s test (**j**). Data are mean ± s.e.m.; \**p* < 0.05, \*\**p* < 0.01, \*\*\**p* < 0.001, \*\*\*\**p* < 0.0001; ns, not significant. See also Extended Data Fig. 11-15.

### Developmental imprints

The long-lived nature of αβ T cells makes the adult landscape a composite of continuous developmental output^61^. Despite transcriptional divergence, the virtual memory layers remain indistinguishable by core markers, precluding retrospective assignment of developmental origin. We used tamoxifen-inducible fate mapping to identify molecular imprints of developmental origin and benchmarked residual *β2m^⁻/⁻^MHCII^⁻/⁻^* CD8⁺ αβ T cells against these imprints to map their ontogeny (Extended Data Fig 14a). The surface glycosphingolipid AsGM1 emerged as the sharpest developmental imprint. Whereas adult-timestamped virtual memory cells largely lacked AsGM1, early-life-timestamped counterparts were predominantly AsGM1^+^ (Fig. 4j). This imprint was recapitulated in humans, with cord-blood virtual memory cells enriched for AsGM1 whereas adult peripheral-blood counterparts were largely AsGM1⁻ (Extended Data Fig 15). Against this developmental reference, *β2m^⁻/⁻^MHCII^⁻/⁻^* cells aligned with the early-life AsGM1⁺ imprint (Fig. 4j,k). CD5 provided an independent quantitative imprint of thymic calibration: expression was lower in early-life-timestamped than adult-timestamped virtual memory cells, with *β2m^⁻/⁻^MHCII^⁻/⁻^* cells again aligning with the early-life state (Extended Data Fig 14b). Because CD5 scales with the strength of thymic TCR calibration^62,63^, this gradient informs the selection environment in which distinct developmental layers of virtual memory were specified. CD122 added a third, complementary axis, revealing an inverse pattern to CD5. This developmental organization gives new context to earlier observations that CD122^lo^ memory-like CD8⁺ αβ T cells are MHC-I- dependent whereas CD122^hi^ cells can be maintained independently of MHC-I^64^ (Extended Data Fig 14b). These imprints resolve the virtual memory compartment into ontogenically distinct branches and independently place the residual *β2m^⁻/⁻^MHCII^⁻/⁻^* population within the early-life route.

## DISCUSSION

Our findings establish that the requirement for MHC in αβ T-cell development is not universal across physiological T-cell fates. Although deficiency of β2m-dependent MHC class I and MHC class II disrupts conventional naive T-cell development^5–8^, the residual αβ compartment does not consist exclusively of stochastic remnants of failed selection. Instead, it resolves into distinct cellular outcomes, including clonally restricted CD4-associated states and a diversified CD8⁺ αβ T-cell population that converges on a physiological wild-type virtual memory state^14,65–67^. This distinction changes the interpretation of a longstanding observation in MHC-deficient mice^5,6,10–12^. The biological significance of the surviving CD8⁺ cells lies not simply in their persistence, but in the alignment of these residual cells with a coherent physiological state across transcriptional and phenotypic identity, as well as TCR repertoire architecture. Their rapid responsiveness to TCR and cytokine stimulation, and their capacity to confer protection during infection, further establish this state as a functionally competent component of host defense.

Developmental timing provides an organizing principle for this alternative outcome. Our data identify the CD8⁺ αβ T-cell fate preserved in MHC-deficient mice as a virtual memory state. However, MHC deficiency does not preserve the full spectrum of physiological virtual memory; instead, it selectively retains states associated with early-life thymopoiesis. Consistent with this developmental restriction, CD8⁺ αβ T cells from adult MHC-deficient mice retain a pronounced neonatal transcriptional imprint and converge phenotypically on the AsGM1⁺ developmental layer identified through early-life timestamping of wild-type virtual memory cells. Fetal progenitors from *β2m^⁻/⁻^MHCII^⁻/⁻^* mice retain the capacity to generate EOMES⁺ CD8⁺ thymocytes without MHC, a capacity largely lost with adult progenitors. The involvement of Lin28b places this developmental asymmetry within a broader principle of heterochronic regulation^57,60,68^. In B lymphopoiesis, Lin28b has been shown to modify the requirements for positive selection during early life, permitting developmental outcomes that are less accessible under adult selection conditions^69–71^. Early-life thymopoiesis may similarly combine a distinct selecting environment with progenitor- intrinsic programs that permit alternative αβ T-cell fates operating under different rules for thymic selection. Notably, the early-life restriction of this alternative fate coincides with a well-established developmental attenuation of the thymic machinery supporting MHC selection. MHC-I expression is low until ten days after birth and MHC-II expression reaches adult levels only after the first two postnatal weeks^72^. Negative selection is also less efficient in newborn mice, as demonstrated by reduced peptide-induced thymocyte deletion^73^, consistent with transcriptional evidence that the thymic machinery supporting central tolerance is developmentally immature during early life ^28,74^. The convergence of these observations places our findings within a physiological developmental context in which alternative requirements for αβ T-cell selection become biologically plausible.

This finding distinguishes the utilization of MHC during physiological development from its absolute requirement for a particular cellular outcome. The preservation of a corresponding virtual memory state in *β2m^⁻/⁻^MHCII^⁻/⁻^* mice demonstrates that the MHC pathways disrupted in this model are not obligatory for every route capable of generating a physiological CD8⁺ αβ T-cell fate. Importantly, the surviving cells lack evidence of conventional self-MHC calibration, arguing against the interpretation that they are normally selected naive T cells that escaped developmental elimination^1,2^. This distinction also separates the generation of an αβ T-cell population from the subsequent acquisition of its mature virtual memory phenotype. Peripheral homeostatic signals can contribute to virtual memory differentiation^38,65,66,75^ but cannot explain how conventionally selected naive CD8⁺ T cells would first arise without the MHC pathways required for their selection^1,2,5,6^. This outcome is conceptually consistent with prior genetic models in which attenuation of TCR signaling redirects αβ T-cell development toward innate-like CD8⁺ fates, suggesting that perturbation of either TCR signaling or MHC calibration can direct αβ T-cell development toward alternative outcomes^24–27,53^. Nonetheless, the present data do not establish that early-life virtual memory cells invariably develop without engaging the residual MHC present in the early-life thymus. What our genetic model establishes is developmental capacity: when MHC selection is unavailable, the early-life thymic environment remains capable of generating a coherent, diversified and functionally competent CD8⁺ αβ virtual memory fate. These findings establish that αβ T-cell specification can proceed under selection conditions that do not generate the conventional naive repertoire, revealing greater developmental flexibility in thymic fate determination.

## ACKNOWLEDGEMENT

We thank Dr. Charles Mainhart and the NIH/NIAID Taconic repository for providing *β2m^⁻/⁻^MHCII^⁻/⁻^* mice. We thank the University of Michigan Platelet Physiology and Pharmacology Core repository and the Program in Research and Innovation on Maternal and Neonatal Outcomes in the Department of Obstetrics and Gynecology for providing de-identified adult peripheral blood and cord blood samples, respectively. We thank the University of Michigan Single Cell Spatial Analysis Shared Resource (RRID:SCR_026842; supported by NIH/NCI P30CA046592), Flow Cytometry Core (RRID:SCR_026697), CyTOF Core Facility (RRID:SCR_026745), ULAM Pathology Core (RRID:SCR_018823), and NIH Tetramer Core Facility (RRID:SCR_026557) for scRNA-seq, flow cytometry, mass cytometry, histopathology, and tetramer reagent support, respectively.

## AUTHOR CONTRIBUTIONS

X.S. performed experiments, analyzed experimental and transcriptomic data, generated figures, performed statistical analyses, and wrote the first manuscript draft. H.G.F. made the initial observation of CXCR3⁺CD8⁺ αβ T-cell persistence in *β2m^⁻/⁻^MHCII^⁻/⁻^* mice. F.M., X.S., A.C.B., L.C.T., and J.E.G. contributed to scRNA-seq clustering and analysis. R.B., L.C., and L.C.T. performed integration analysis of IL-4-induced CXCR3^+^CD8⁺ αβ T-cell datasets. X.S. performed downstream interpretation and cross-referencing of scRNA-seq datasets with public transcriptional references. A.J.Z., X.S., F.W., and Y.L. designed CyTOF experiments and antibody panel. S.M.R., A.J.Z., I.O.V., and F.W. performed CyTOF technical optimization, acquisition, and data analysis. L.O.C.-M. and S.A.C. generated the LCMV infection–derived memory CD8⁺ αβ T-cell model. X.S. performed FTOC and RTOC experiments under S.A.C.’s guidance. P.T.R., C.S., and S.M.M.H. performed MR1 tetramer–based analyses to assess MAIT cell representation. K.A.E. performed histopathological evaluation of tissue sections. X.S., F.M., A.J.Z., R.B., A.C.B., L.C.T., J.E.G., F.W., P.T.R., S.M.M.H., K.A.E., A.L., and Y.L. contributed to data visualization and figure design. A.L. contributed to conceptual development, organization of the data narrative, manuscript writing and editing, infection-related experiments, and key reagents. Y.L. was responsible for overall conceptualization, funding acquisition, project administration, supervision, data interpretation, and manuscript writing.

## Extended data figures

**Extended Data Fig. 1:**
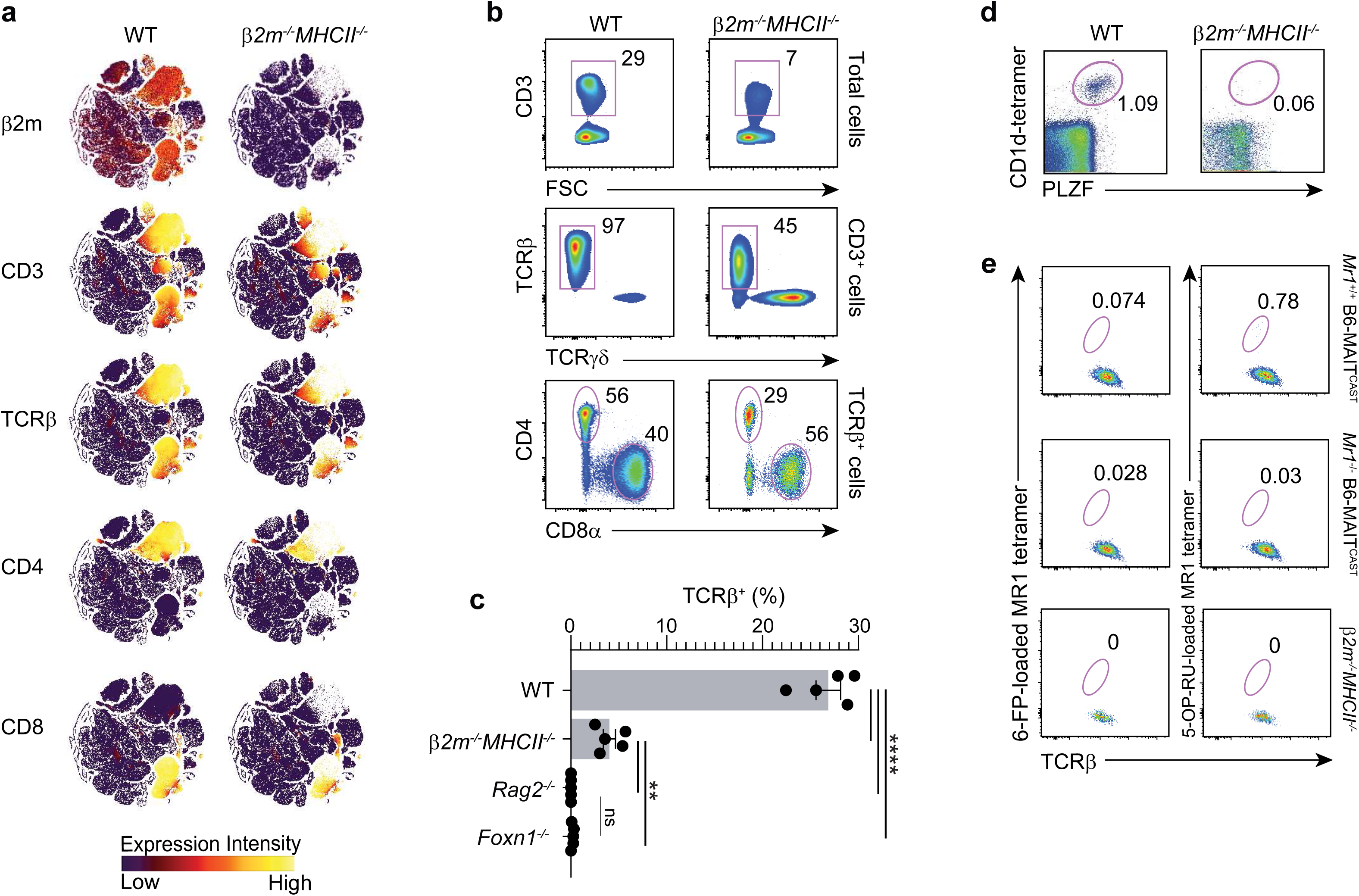
Peripheral αβ T-cell composition in *β2m^⁻/⁻^MHCII^⁻/⁻^ mice*. **a,** Relative expression of indicated markers across the immune landscape from CyTOF analysis in Fig 1a. **b,** Representative gating strategy to represent the frequency of CD3ε^+^ among splenocytes, TCRβ^+^ among CD3ε^+^ splenocytes, CD4/CD8α distribution among CD3ε^+^TCRβ^+^ splenocytes from WT and *β2m^-/-^MHCII^-/-^* mice. **c,** Frequency of splenic CD3⁺TCRβ⁺ cells in WT, *β2m^-/-^MHCII^-/-^*, *Rag2^⁻/⁻^*, and *Foxn1^⁻/⁻^* mice. **d,** Distribution of PLZF⁺CD1d tetramer⁺ cells among splenic TCRβ⁺ cells from WT and *β2m^-/-^MHCII^-/-^* mice. **e,** MR1 tetramer staining of splenic TCRβ⁺CXCR3⁺ cells from *Mr1*^+/+^ and *Mr1*^-/-^ B6-MAIT^CAST^and *β2m^-/-^MHCII^-/-^* mice. *Mr1*^+/+^ and *Mr1*^- /-^ B6-MAIT^CAST^ mice were initially provided by Dr. Olivier Lantz (Insitut Curie, Paris, France), bred and maintained at Western University (London, Ontario, Canada) and used as positive and negative controls, respectively. 6-FP-loaded mouse MR1 tetramer was used as a negative staining control to set the respective cytofluorimetric gate. Data (**b-e**) are from 2-6 independent experiments; n = 2–14. One-way ANOVA with Tukey’s test (**c**). Data are mean ± s.e.m.; \**p* < 0.05, \*\**p* < 0.01, \*\*\**p* < 0.001, \*\*\*\**p* < 0.0001; ns, not significant.

**Extended Data Fig. 2:**
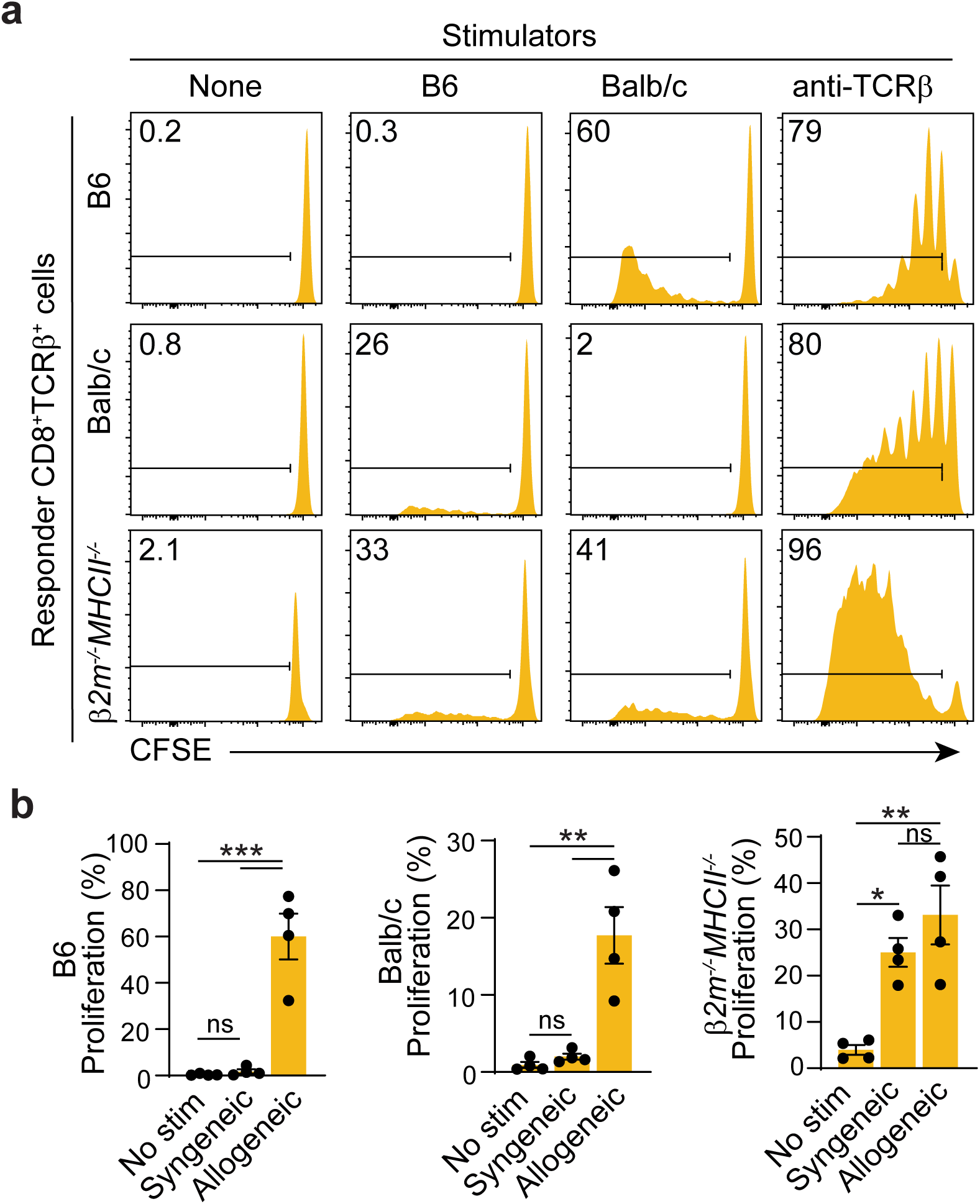
CD8⁺ αβ T cells from *β2m^⁻/⁻^MHCII^⁻/⁻^* mice are MHC-uncalibrated. CFSE-labeled splenic αβ T-cell responders were cultured for 4 days without stimulators, with B6 or BALB/c splenocyte stimulators, or with anti-TCRβ (2 µg/ml). Representative CFSE dilution in responder CD8⁺ αβ T cells (**a**) and frequency of dividing responder CD8⁺ αβ T cells (**b**) are shown. Data are from 4 independent experiments; n = 4. One-way ANOVA with Holm-Šídák’s test. Data are mean ± s.e.m.; \**p* < 0.05, \*\**p* < 0.01, \*\*\**p* < 0.001, \*\*\*\**p* < 0.0001; ns, not significant.

**Extended Data Fig. 3:**
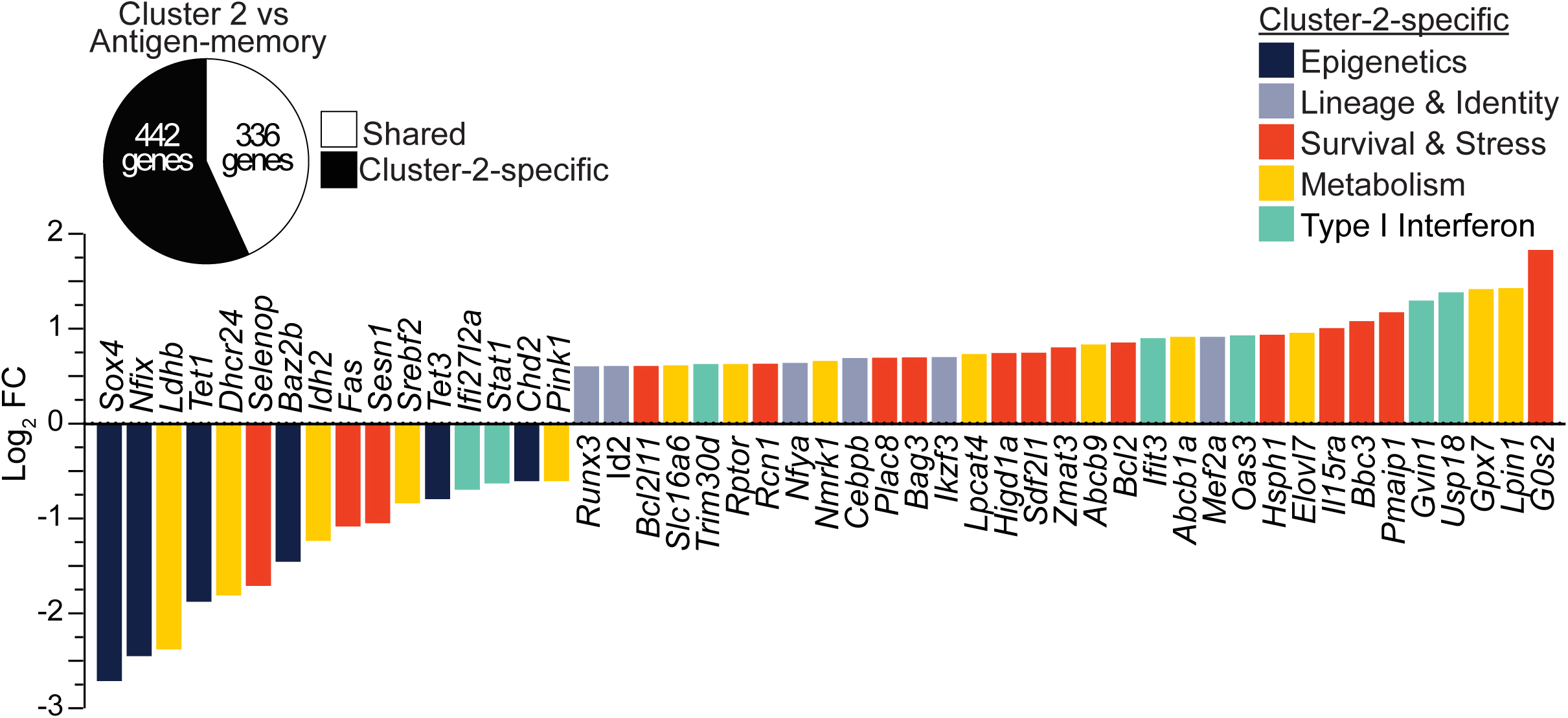
*β2m^⁻/⁻^MHCII^⁻/⁻^* CD8⁺ αβ T cells are transcriptionally distinct from antigen-driven memory. Cross-dataset comparison of cluster 2 vs. cluster 1 DEGs in Fig 2a against reference antigen- memory vs. naive CD8⁺ T-cell DEGs (GSE112304; antigen-memory, CD3⁺CD8⁺CD44⁺CD49d⁺; naive, CD3⁺CD8⁺CD44⁻)^42^. Pie chart stratifying the 778 cluster 2 vs. cluster 1 DEGs into antigen- memory-shared and cluster-2-specific genes (top). Selected cluster-2-specific DEGs grouped by biological function (bottom). DEGs were defined as log_2_ fold change > 0.6 or < −0.6, adjusted *p* < 0.05.

**Extended Data Fig. 4:**
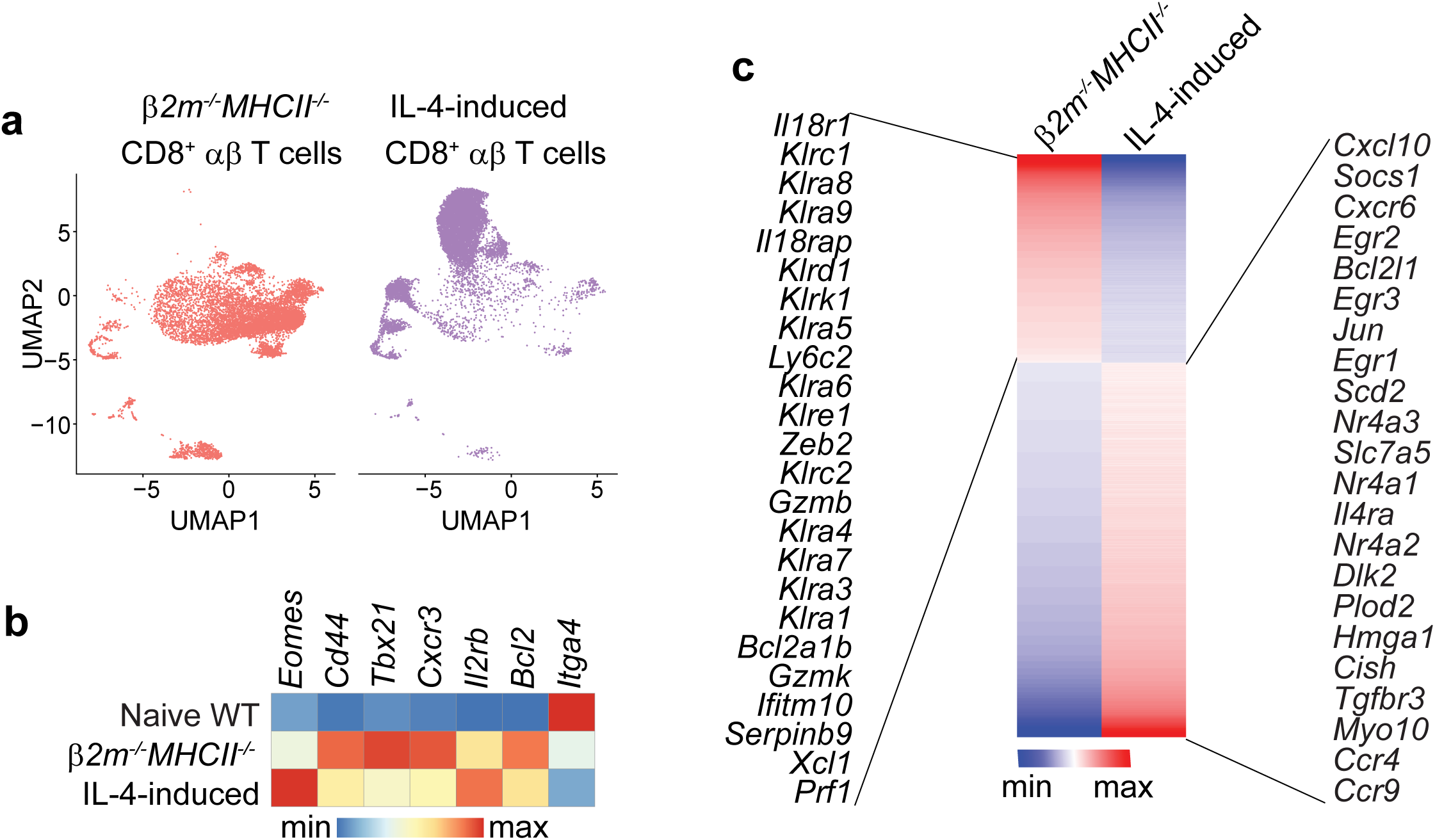
*β2m^⁻/⁻^MHCII^⁻/⁻^* CD8⁺ αβ T cells are transcriptionally distinct from IL- 4-induced memory-like T cells. Integrated scRNA-seq analysis comparing FACS-sorted splenic WT CXCR3^-^CD8^+^ αβ T cells, WT CXCR3^+^CD8^+^ αβ T cells, *β2m^-/-^MHCII^-/-^*CXCR3^+^CD8^+^ αβ T cells and WT IL-4-induced CXCR3⁺CD8⁺ αβ T cells. **a,** UMAP distribution of *β2m^-/-^MHCII^-/-^* CXCR3^+^CD8^+^ αβ T cells and WT IL-4-induced CXCR3⁺CD8⁺ αβ T cells. **b,** Virtual memory-associated gene expression. **c,** heatmap of 2713 DEGs identified from *β2m^-/-^MHCII^-/-^*CXCR3^+^CD8^+^ αβ T cells over WT IL-4- induced CXCR3⁺CD8⁺ αβ T cells; selected genes are labeled. DEGs were defined as log_2_ fold change > 0.6 or < −0.6, adjusted *p* < 0.05.

**Extended Data Fig. 5:**
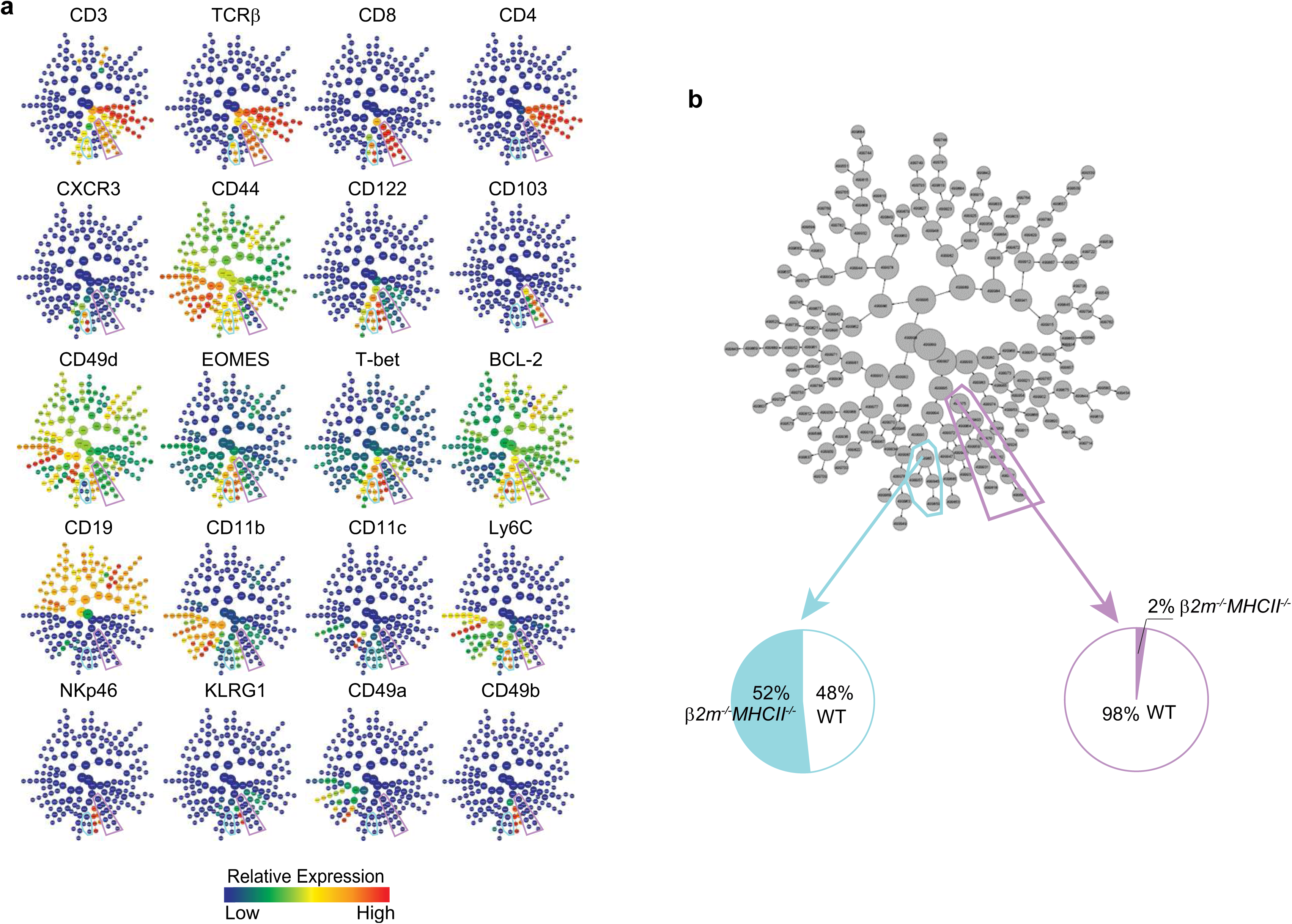
CITRUS identifies virtual memory as the predominant residual CD8⁺ αβ T-cell state in *β2m^⁻/⁻^MHCII^⁻/⁻^* mice. **a,** CITRUS clustering of splenocytes from CyTOF analysis in Fig 1a with relative expression of indicated markers; purple and blue gates mark naive and virtual memory branches, respectively. **b,** WT and *β2m^⁻/⁻^MHCII^⁻/⁻^* sample distribution within naive and virtual memory parent clusters. Data are from n = 5 per genotype.

**Extended Data Fig. 6:**
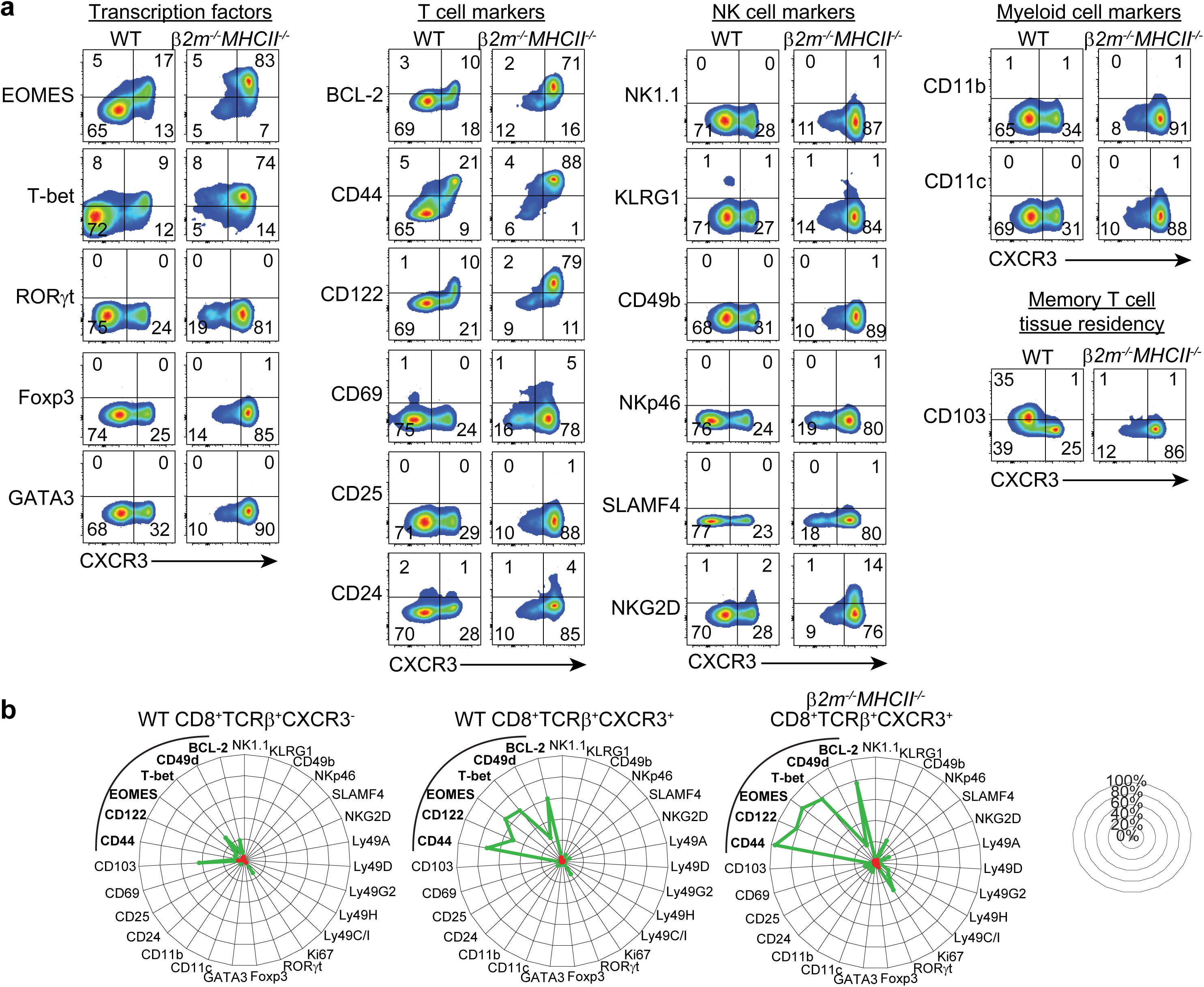
Comprehensive phenotypic profiling of the residual CD8⁺ αβ T-cell compartment in *β2m^⁻/⁻^MHCII^⁻/⁻^* mice. **a,** Distribution of CXCR3 versus each indicated marker among splenic CD8⁺TCRβ⁺ cells from WT and *β2m^-/-^MHCII^-/-^*mice. Plots are grouped by transcription factors, T cell markers, NK cell markers, myeloid cell markers, and memory T cell tissue-residency markers. **b,** Radar plots of marker-expression frequency among the indicated CD8⁺ αβ T-cell subsets. Green lines indicate mean frequency and red lines indicate s.e.m.; concentric circles indicate 20% frequency increments. Data are from 2-6 independent experiments; n = 4-18.

**Extended Data Fig. 7:**
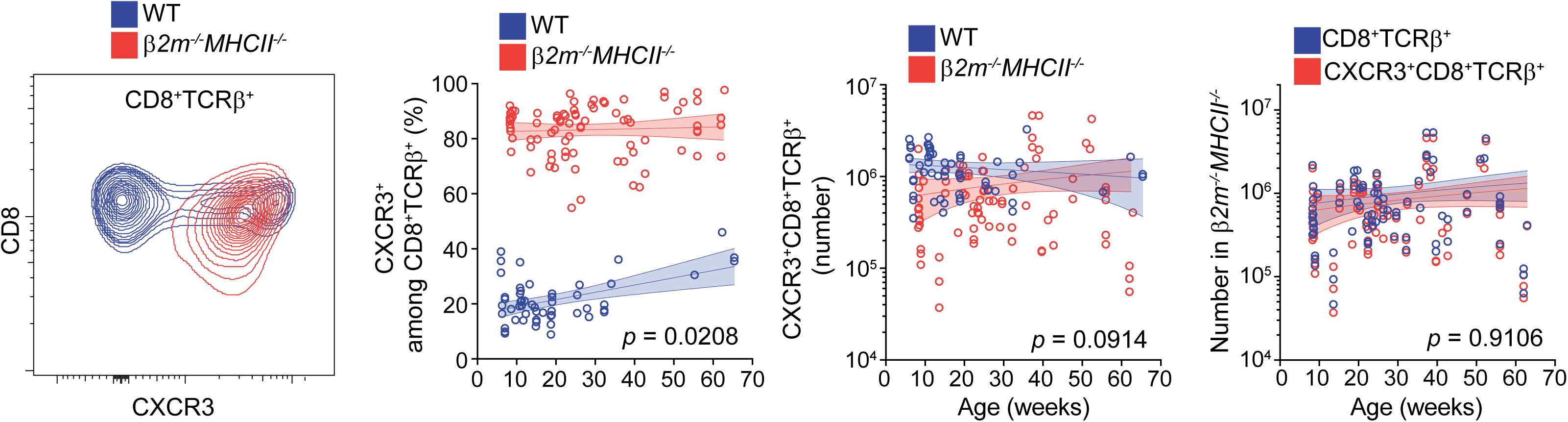
Age-dependent dynamics of the CD8⁺ αβ T-cell compartment in *β2m^⁻/⁻^MHCII^⁻/⁻^* mice. Representative CXCR3 expression among splenic CD8⁺ αβ T cells in WT and *β2m^⁻/⁻^MHCII^⁻/⁻^* mice (flow cytometry plot). Frequency and number of CD8⁺ αβ T-cell subsets across age (kinetic dot plots). Data (flow cytometry plot) are from 2-6 independent experiments; n = 11-14. Kinetic dot plots include n = 54–84 mice across the indicated age range. For kinetic dot plots, lines indicate simple linear regression with 95% confidence bands. P values indicate ANCOVA comparisons of slopes between groups.

**Extended Data Fig. 8:**
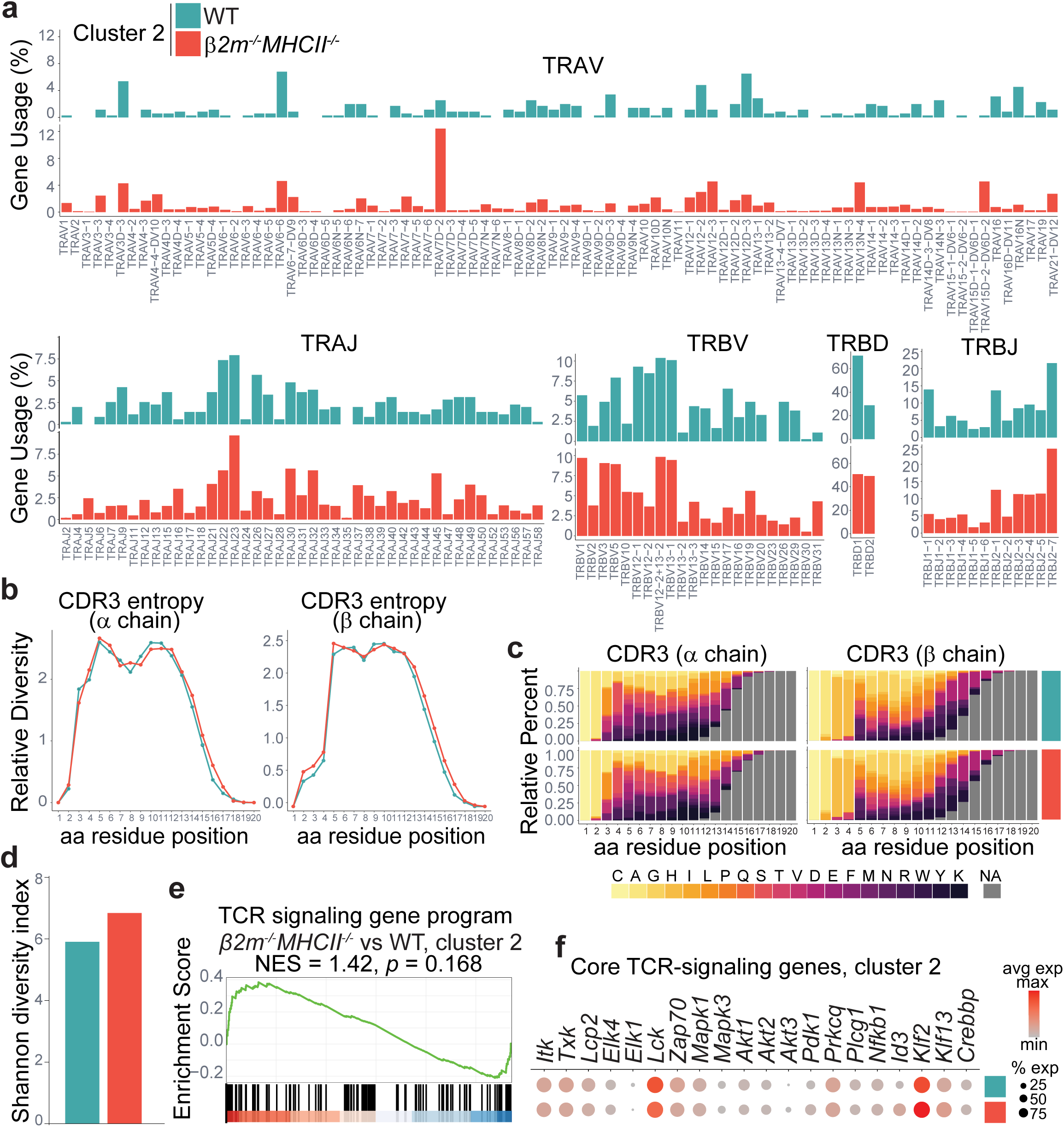
TCR diversity is preserved in MHC-resilient CD8⁺ αβ T cells. **a-d,** scTCR immune profiling of defined CD8⁺ αβ T-cell populations: cluster 2 WT and cluster 2 *β2m^⁻/⁻^MHCII^⁻/⁻^* cells in Fig 2a. **a,** TCRα V/J and TCRβ V/D/J gene usage. **b,** CDR3 positional amino-acid (aa) entropy across positions 1–20 for TCRα and TCRβ chains. **c,** Distribution of aa frequencies across CDR3 positions 1–20 for TCRα and TCRβ chains. **d,** Shannon diversity index of the TCR clone repertoire. **e,f,** TCR signaling analysis in cluster 2 cells, comparing *β2m^⁻/⁻^MHCII^⁻/⁻^* and WT cells. **e,** GSEA of the TCR signaling pathway. NES, normalized enrichment score. **f,** Dot plot of indicated TCR signaling genes.

**Extended Data Fig. 9:**
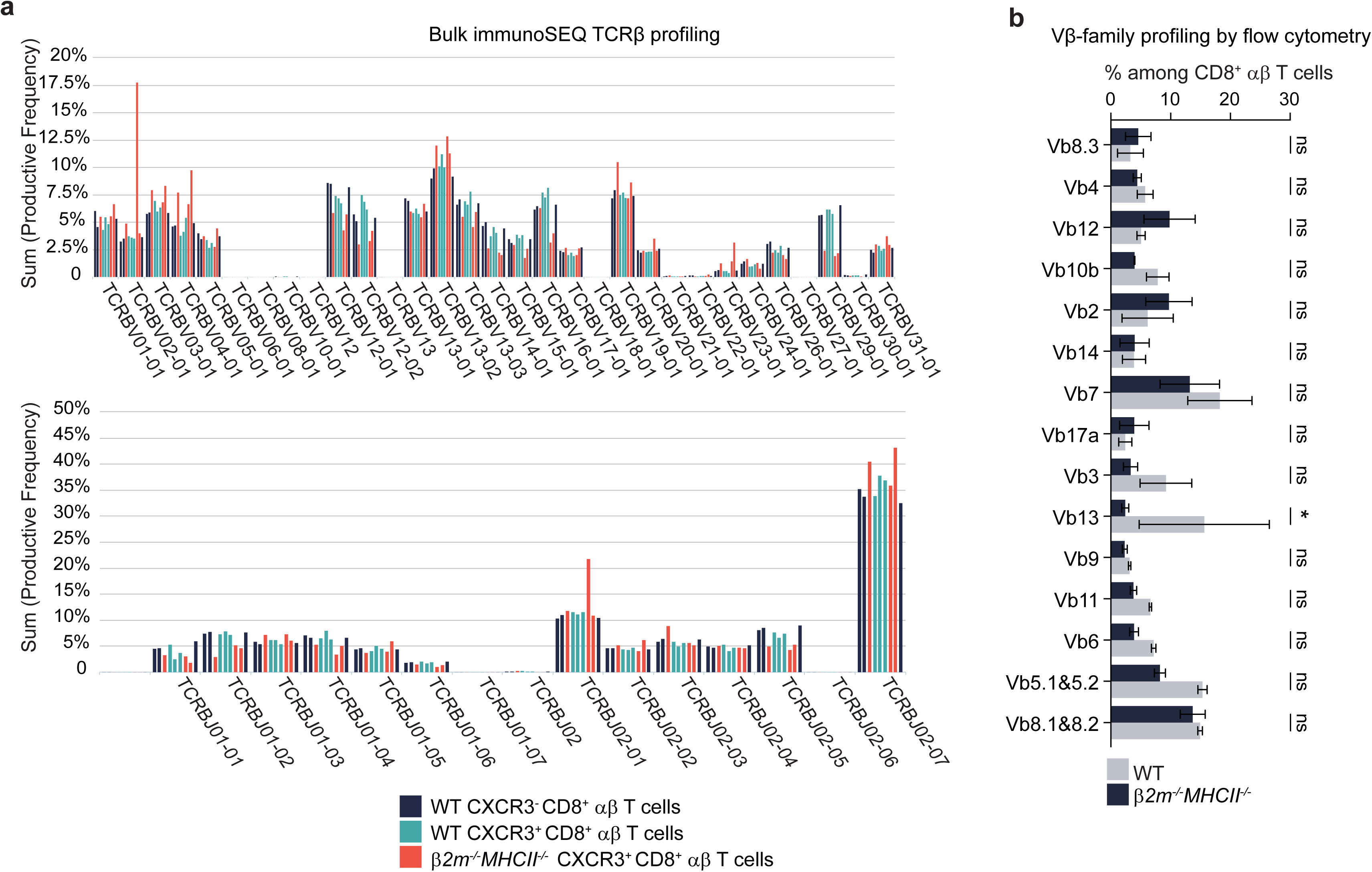
TCRβ gene usage is preserved in CD8⁺ αβ T cells from *β2m^⁻/⁻^MHCII^⁻/⁻^* mice. **a,** ImmunoSEQ analysis of TCRBV (top) and TCRBJ (bottom) gene usage, quantified as summed productive frequency, in FACS-sorted WT CXCR3⁻CD8⁺ αβ T cells, WT CXCR3⁺CD8⁺ αβ T cells, and *β2m^-/-^MHCII^-/-^* CXCR3⁺CD8⁺ αβ T cells. Data are shown for triplicate samples. **b,** Flow- cytometric quantification of TCR Vβ-chain–positive cells among splenic CD8⁺ αβ T cells from WT and *β2m^-/-^MHCII^-/-^* mice. Data (**b**) are from 2 independent experiments; n = 3-5. Two-way ANOVA with Šídák’s test. Data are mean ± s.e.m.; \**p* < 0.05, \*\**p* < 0.01, \*\*\**p* < 0.001, \*\*\*\**p* < 0.0001; ns, not significant.

**Extended Data Fig. 10:**
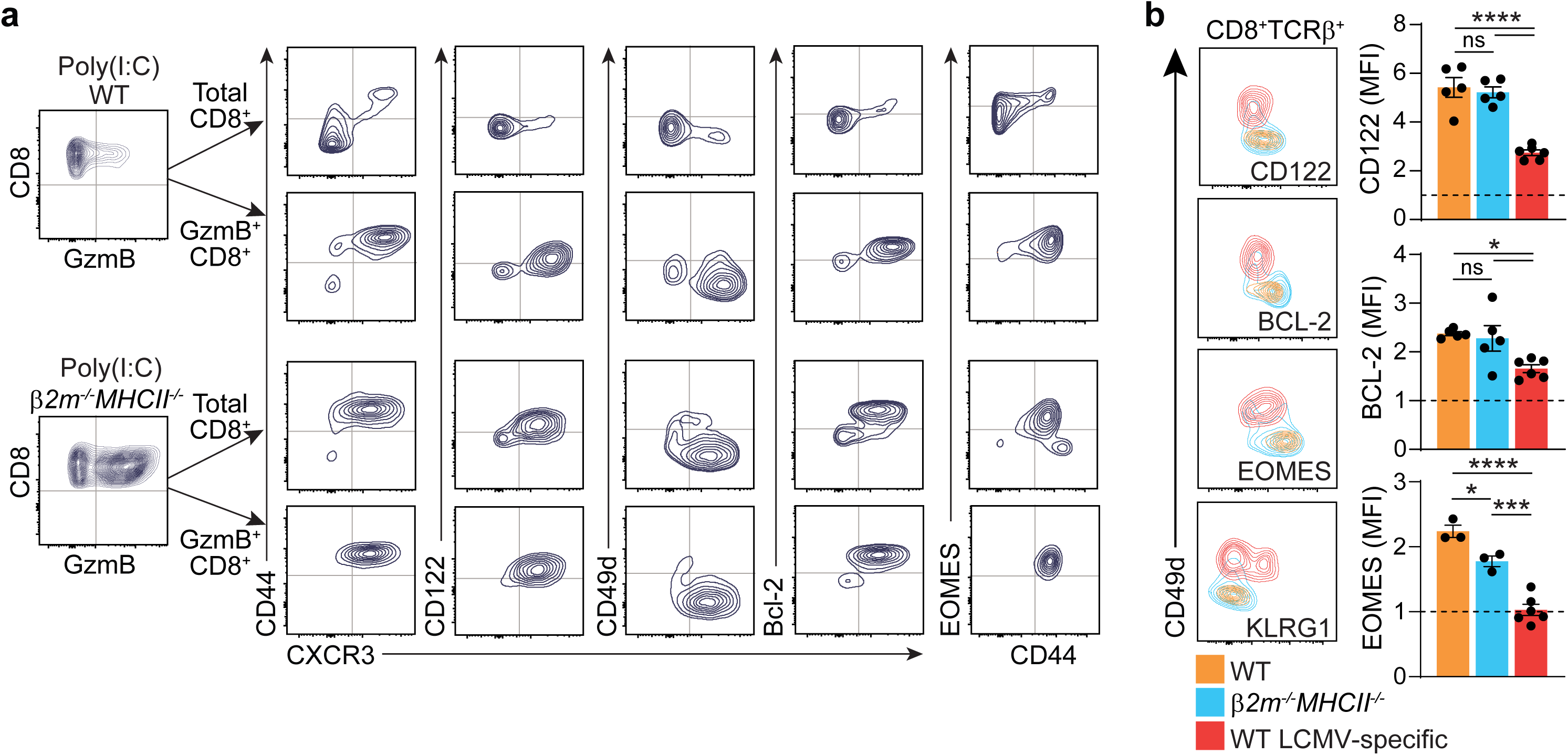
Inflammatory responsiveness of *β2m^⁻/⁻^MHCII^⁻/⁻^* CD8⁺ αβ T cells. **a,** GzmB expression among CD8⁺ αβ T cells from WT and *β2m^⁻/⁻^MHCII^⁻/⁻^* mice, with marker distribution in total vs. GzmB⁺ CD8⁺ αβ T cells. **b,** Overlay of WT CD44^+^CD49d^-^ CD8⁺ αβ T cells, *β2m^⁻/⁻^MHCII^⁻/⁻^* CD8⁺ αβ T cells, and NP396 tetramer⁺ LCMV-specific memory CD8⁺ αβ T cells (CD44⁺CD49d⁺) on CD49d vs. indicated marker plots. Marker-expression fold change in each subset relative to naive WT CD8⁺ αβ T cells. MFI, mean fluorescence intensity. Data are from 2- 3 independent experiments; n = 3-8 mice. One-way ANOVA with Tukey’s test (**b**). Data are mean ± s.e.m.; \**p* < 0.05, \*\**p* < 0.01, \*\*\**p* < 0.001, \*\*\*\**p* < 0.0001; ns, not significant.

**Extended Data Fig. 11:**
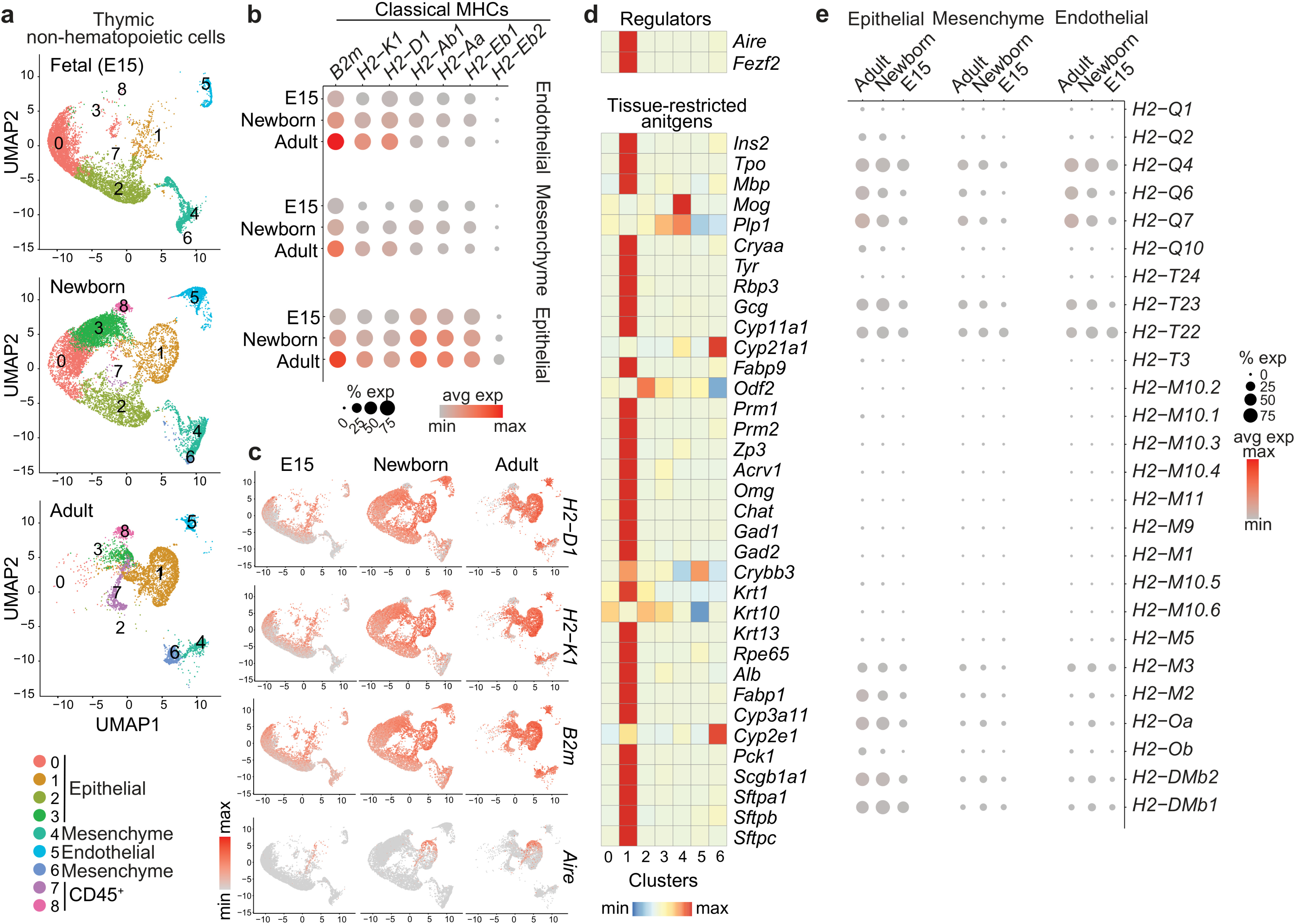
Thymic MHC selection machinery is developmentally staged. scRNA-seq data from thymic non-hematopoietic cells isolated from fetal, newborn, and adult WT mice (GSE192480)^28^. **a,** UMAP distribution by developmental stage; clusters are annotated as epithelial, mesenchymal, or endothelial, with residual CD45⁺ hematopoietic contaminants indicated. **b,** Dot plots of classical MHC class I and class II genes across stromal compartments and developmental stages. **c,** Feature plots of indicated genes across developmental stages. **d,** Heatmap of central tolerance regulators and tissue-restricted antigen genes across clusters. **e,** Dot plot of nonclassical MHC class I and MHC class II accessory gene expression across age groups and stromal compartments.

**Extended Data Fig. 12:**
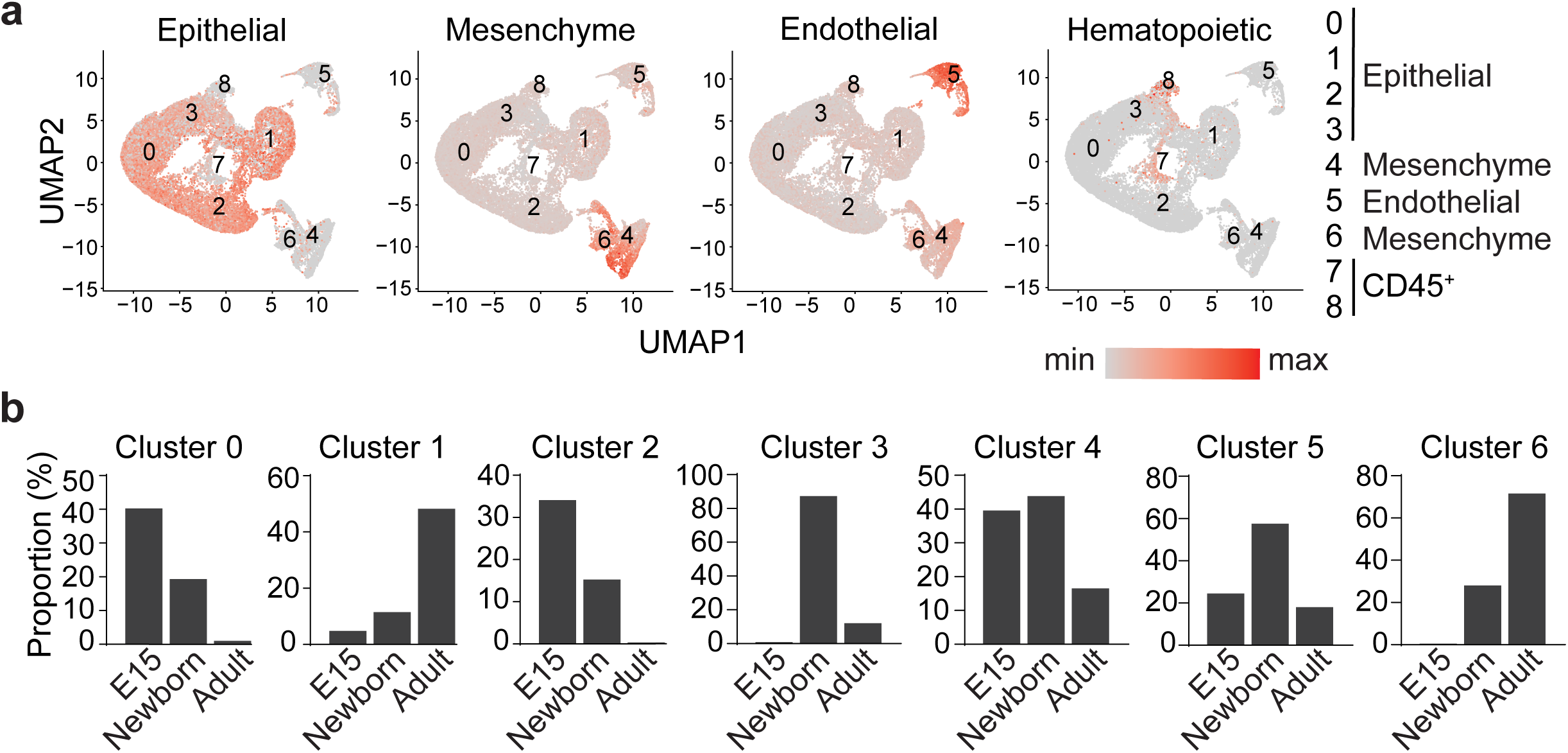
Ontogeny of thymic stromal compartments. Analysis of a public thymic stromal (CD45^-^EpCAM^+^) scRNA-seq dataset (GSE192480)^28^ from E15 fetal, newborn, and adult WT mice. **a,** Feature plots showing epithelial, mesenchymal, endothelial clusters annotated using epithelial (*Epcam*), mesenchymal (*Col3a1*, *Pdgfra*, *Fbn1*, *Dlk1*, *Sfrp2*, *Gdf10*, *Igf1*, *Igf2*), and endothelial (*Pecam1*, *Cdh5*, *Eng*, *Cldn5*, *Cd36*, *Egfl7*, *Igfbp7*, *Sox17*) markers. Residual CD45⁺ hematopoietic contaminants were identified by *Ptprc* expression. **b,** Age-group composition of each cluster.

**Extended Data Fig. 13:**
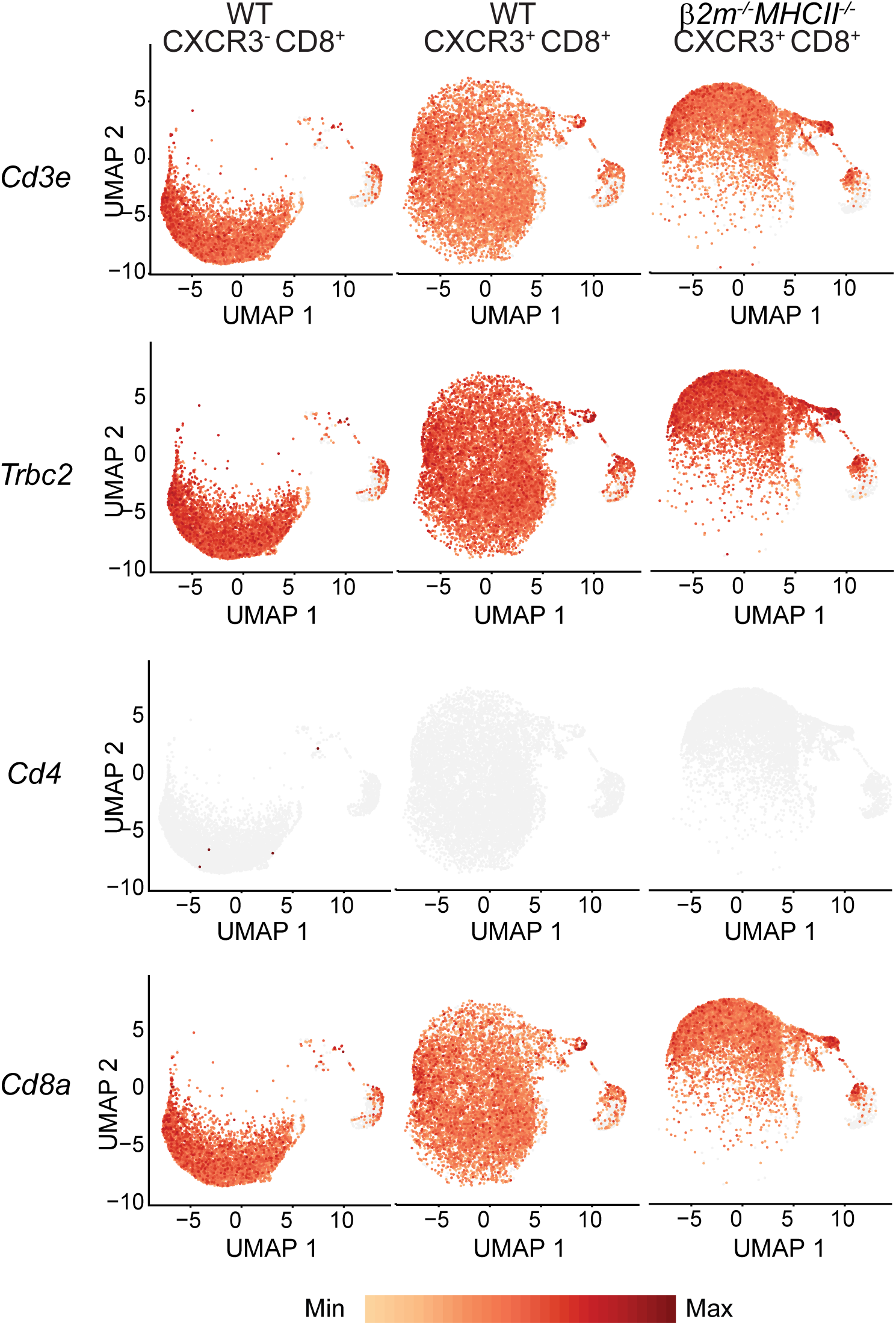
scRNA-seq annotation of sorted CD8⁺ αβ T-cell populations. Feature plots showing *Cd3e*, *Trbc2*, *Cd4*, and *Cd8a* expression across scRNA-seq datasets from FACS-sorted splenic WT CXCR3^-^CD8^+^ αβ T cells, WT CXCR3^+^CD8^+^ αβ T cells, and *β2m^-/-^MHCII^- /-^* CXCR3^+^CD8^+^ αβ T cells, gated as CD3⁺TCRγδ⁻TCRβ⁺CD8⁺ cells. Gene expression confirms CD8 αβ T-cell annotation across datasets.

**Extended Data Fig. 14:**
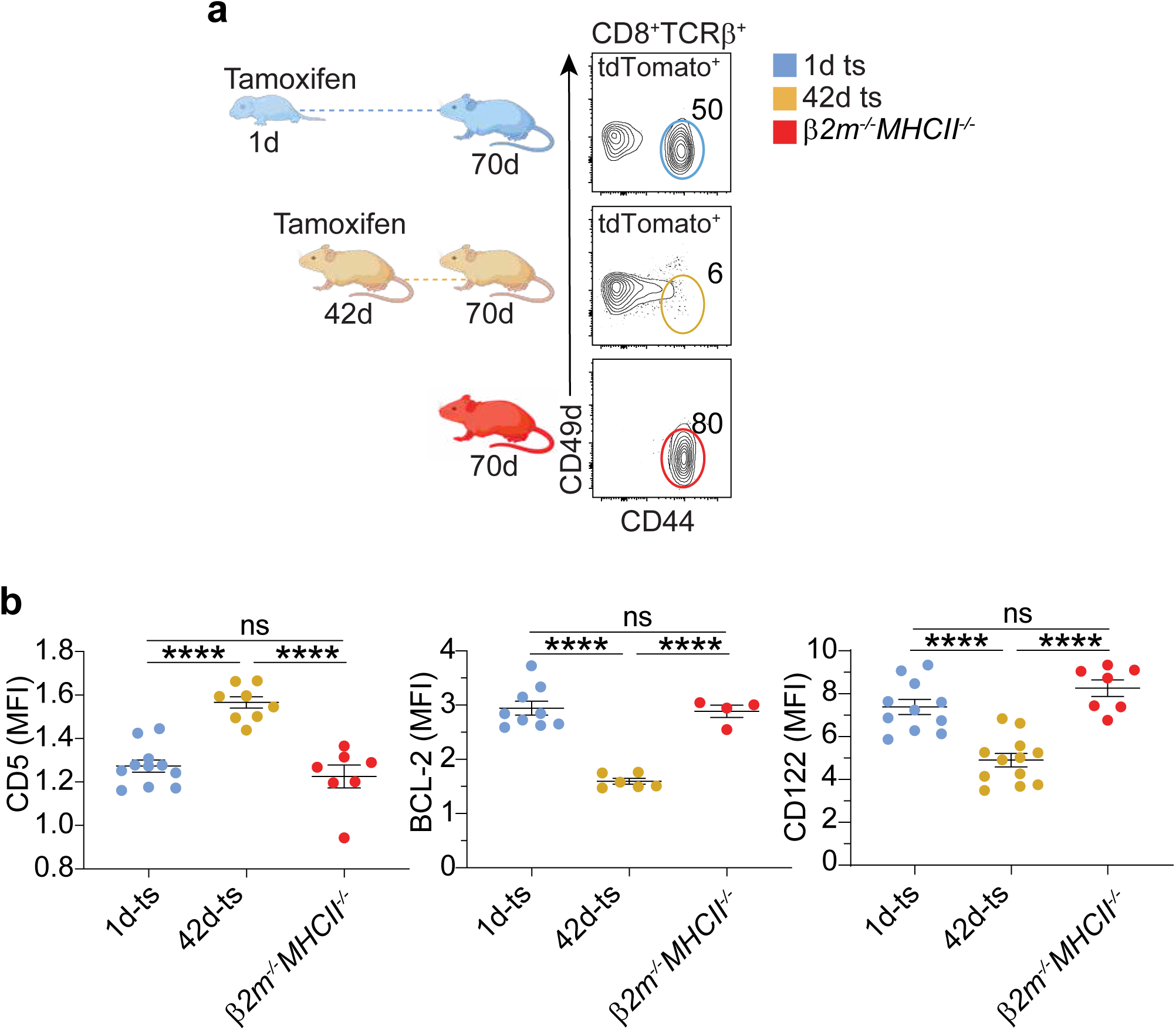
Developmental imprints of virtual memory T-cell origin revealed by mouse timestamping. **a**, Timestamping scheme and CD44/CD49d distribution among splenic CD8⁺ αβ T cells from *β2m^⁻/⁻^MHCII^⁻/⁻^* mice and tdTomato⁺ CD8⁺ αβ T cells from 1-day timestamped (1d ts) and 42-day timestamped (42d ts) mice. Gates and frequencies indicate virtual memory T cells. **b**, CD5, BCL- 2, and CD122 expression among indicated virtual memory T cell populations. MFI values are normalized to naive WT CD8⁺ αβ T cells. Data (**b**) are from 2-6 independent experiments; n = 4-12. One-way ANOVA with Tukey’s test (**b**). Data are mean ± s.e.m.; \**p* < 0.05, \*\**p* < 0.01, \*\*\**p* < 0.001, \*\*\*\**p* < 0.0001; ns, not significant.

**Extended Data Fig. 15:**
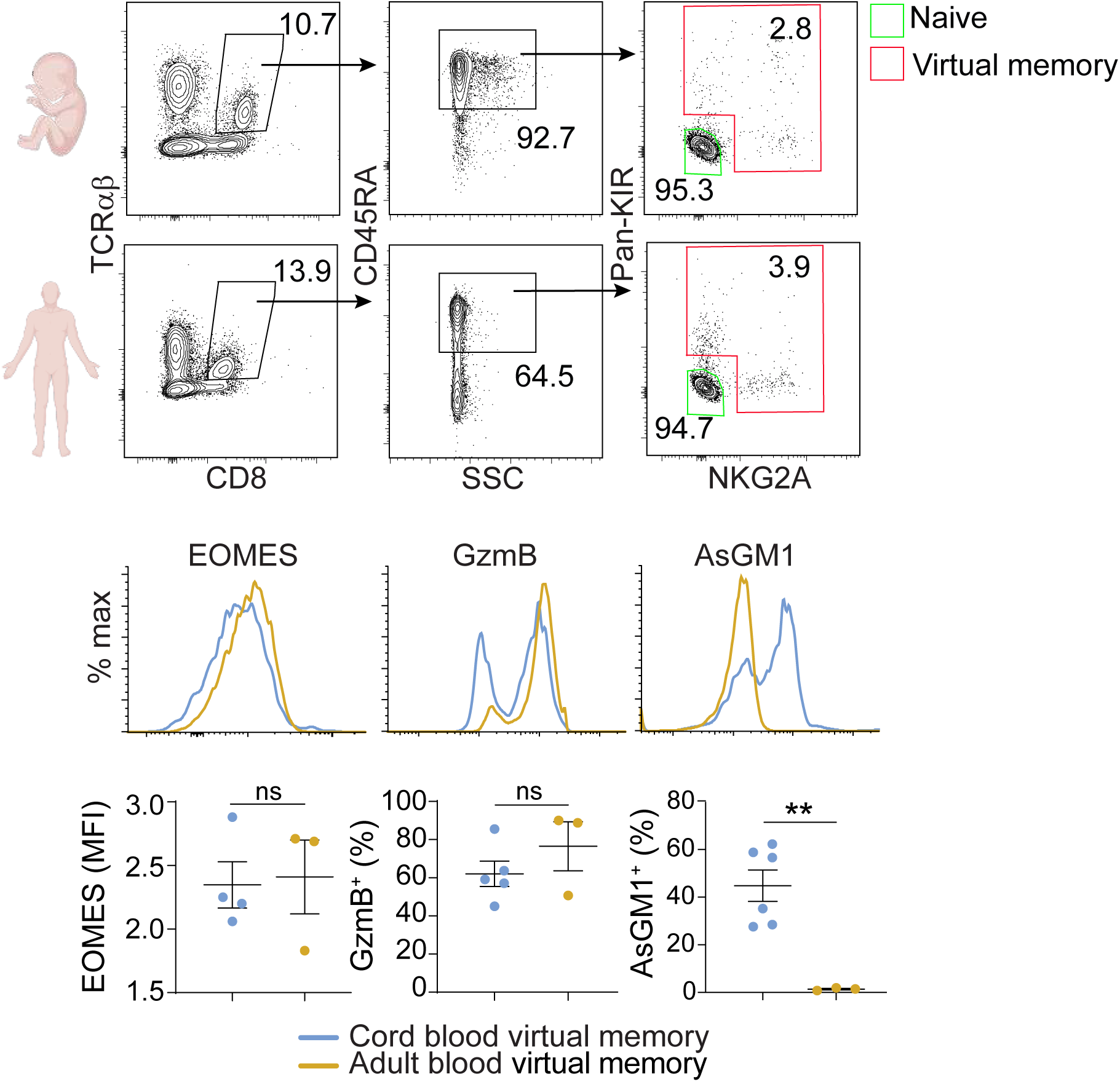
AsGM1 marks an early-life virtual memory T-cell imprint conserved in humans. Flow cytometric analysis of human virtual memory T cells (CD8⁺TCRαβ⁺CD45RA⁺ expressing Pan-KIR and/or NKG2A)^76,77^ from cord blood and adult peripheral blood. Representative gating strategies for human naive and virtual memory T cells (top). EOMES, GzmB, and AsGM1 expression of virtual memory T cells in human cord blood vs. adult peripheral blood shown by histograms; GzmB was assessed 24 h after IL-15 stimulation (50 ng/ml) (middle). GzmB^+^ and AsGM1⁺ frequency and EOMES expression among indicated virtual memory T cell populations. MFI values are normalized to cord blood naive CD8⁺ αβ T cells (bottom). Data are from 2-3 independent experiments; n = 3-6. unpaired two-tailed t-test. Data are mean ± s.e.m.; \**p* < 0.05, \*\**p* < 0.01, \*\*\**p* < 0.001, \*\*\*\**p* < 0.0001; ns, not significant.

## METHODS

### Mice

B6 (C57BL/6J) (#000664), CD45.1 B6 (B6.SJL-*Ptprc^a^ Pepc^b^*/BoyJ) (#002014), BALB/c (BALB/cJ) (#000651), CD45.1 BALB/c (CByJ.SJL(B6)-*Ptprc^a^*/J) (#006584), CD4CreER^T2^ (B6(129X1)-Tg(Cd4-cre/ERT2)11Gnri/J) (#022356), Ai9 (B6.Cg-*Gt(ROSA)26Sor^tm9(CAG-tdTomato)Hze^*/J) (#007909), *Rag2^-/-^* (B6.Cg-*Rag2^tm1.1Cgn^*/J) (#008449), and nude *Foxn1^⁻/⁻^* (#002019) mice were purchased from The Jackson Laboratory. *Rag2^⁻/⁻^Il2rγ^⁻/⁻^* mice (C57BL/6NTac.Cg- *Rag2^tm1Fwa^ Il2rg^tm1Wjl^*) (#4111) were purchased from Taconic Biosciences. *Mr1*^+/+^ and *Mr1*^-/-^ B6- MAIT^CAST^ mice were provided by Dr. Olivier Lantz (Institut Curie, Paris, France). *β2m^⁻/⁻^MHCII^⁻/⁻^* mice (B6.129-*H2-Ab1^tm1Gru^ B2m^tm1Jae^* N17) (#4080) were provided by the NIH/NIAID Taconic repository. CD45.1 *β2m^⁻/⁻^MHCII^⁻/⁻^* mice were generated by crossing CD45.1 B6 mice with *β2m^⁻/⁻^MHCII^⁻/⁻^* mice. Wild-type (WT) control mice are either B6 or CD45.1 B6. Male and female mice from birth to 65 weeks of age were used, and embryos were collected at embryonic day 15.5. Age-matched B6 mice served as wild-type (WT) controls. All mice were maintained under specific pathogen-free conditions at the University of Michigan, and all procedures were approved by the University of Michigan Institutional Animal Care and Use Committee (protocols PRO00012457 and PRO00010912) in accordance with the Guide for the Care and Use of Laboratory Animals.

### Human Samples

De-identified adult peripheral blood samples were obtained from the University of Michigan Platelet Physiology and Pharmacology Core repository. De-identified human cord blood samples were obtained through the Program in Research & Innovation on Maternal Outcomes with approval from the Obstetrics and Gynecology Labor and Delivery Tissue Committee at Von Voigtlander Women’s Hospital under University of Michigan IRB-approved protocol HUM00110708.

### Bacterial and viral strains

Listeria monocytogenes 10304S was obtained from laboratory stocks maintained by Dr. Amale Laouar laboratory. LCMV Armstrong was obtained from laboratory stocks maintained by Dr. Shannon Carty.

### Poly(I:C) treatment of neonatal and adult mice

Poly(I:C) (#P1530-100MG, Sigma-Aldrich) was administered intraperitoneally at 3.75 mg/kg body weight to either adult (6–15 weeks old) or neonatal (1 week old) mice. Mice were weighed before injection to determine dosing. Approximately 18 h after treatment, mice were euthanized, and splenocytes were harvested for analysis of intracellular granzyme B expression.

### Listeria monocytogenes infection

Listeria monocytogenes (LM), provided by Dr. Amale Laouar (University of Illinois Urbana- Champaign), was cultured in brain-heart infusion (BHI) medium (#53286-100G, Sigma-Aldrich). Infectious doses were determined using an OD–CFU standard curve generated from hourly measurements of bacterial growth. Mice were infected intraperitoneally with 10^5^ CFU LM and splenocytes were harvested 48 h post-infection for analysis of IFNγ expression. For adoptive- transfer LM infection experiments, donor splenic CD8^+^ αβ T cells were enriched by negative depletion of TCRγδ⁺ (#118103, BioLegend), IgD⁺ (#405733, BioLegend), IgM⁺ (#406503, BioLegend), CD4⁺ (#13004282, eBioscience), NK1.1⁺ (#108703, BioLegend), and CD11b⁺ (#101203, BioLegend) cells using the EasySep™ Mouse Streptavidin RapidSpheres Isolation Kit (#19860, STEMCELL Technologies), followed by FACS sorting into TCRβ⁺CD8α⁺CXCR3⁻ and TCRβ⁺CD8α⁺CXCR3⁺ populations. *Rag2^⁻/⁻^Il2rγ^⁻/⁻^* recipient mice received 0.2 x 10^6^ sorted donor cells intravenously 24 h before intravenous challenge with 10^4^ CFU LM. Bacterial burden was assessed 72 h post-infection by plating serial dilutions of homogenized spleen and liver tissues on BHI agar plates and calculating CFU per gram of tissue.

### Lymphocytic choriomeningitis virus (LCMV)-Armstrong infection

Mice were infected intraperitoneally with 2 x 10^5^ PFU of LCMV Armstrong strain. At day 45 post- infection, mice were treated with poly(I:C) as described above. LCMV-specific memory CD8^+^ αβ T cells were identified using NP396 tetramer (NIH Tetramer Core Facility) staining for subsequent analysis.

### Pharmacologic inhibition of Lin28b in neonatal mice

Lactating dams received daily intravenous injections of C1632 (250 µg in PBS (#BP6651, FISHER scientific) with 5% DMSO (#D5879-500ML, Sigma-Aldrich); #SML3673, Sigma-Aldrich)^78^ for 14 days following parturition. Neonatal mice were exposed to C1632 indirectly through maternal lactation. Pups were euthanized at postnatal day 7 for thymocyte analysis and at postnatal day 14 for splenocyte analysis.

### *In vivo* IL-4C induction of CXCR3^+^CD8^+^ αβ T cells

WT mice received daily intraperitoneal injections of IL-4 complex (IL-4C) consisting of 1.5 µg recombinant murine IL-4 (#214-14, PeproTech) and 50 µg anti–mouse IL-4 antibody (#BE0045, Bio X Cell) for 7 days, as previously described^52^. Splenic IL-4-induced CXCR3^+^CD8^+^ αβ T cells were flow sorted as CD3^+^TCRβ^+^TCRγδ^-^CD4^-^CD8^+^CXCR3^+^CD44^+^CD124^+^ cells and used for single-cell RNA sequencing.

### Tamoxifen-induced CD8⁺ αβ T-cell timestamping

CD4CreERT2 mice were crossed with Ai9 reporter mice to generate CD4CreERT2 ^+/−^Ai9^+/−^ experimental mice, with Cre-negative Ai9^+/−^ littermates used as controls. To induce tdTomato expression, tamoxifen (#T5648, Sigma-Aldrich) dissolved in peanut oil (#P2144, Sigma-Aldrich) was administered via oral gavage. For neonatal timestamping, lactating dams received three 2.5 mg tamoxifen doses between postpartum day 0 and day 1, allowing indirect tamoxifen transfer to pups through milk. For adult timestamping, 42-day-old mice received two 2 mg tamoxifen doses over a 24 h period. Mice were analyzed at 70 days of age.

### Tissue processing and single-cell preparation

Single-cell suspensions from spleen were prepared in PBS supplemented with 5% fetal bovine serum (FBS; #F4135, Sigma-Aldrich). Erythrocytes were lysed with red blood cell lysis buffer containing 170 mM Tris and 160 mM NH₄Cl in Milli-Q water. Cells were filtered through 40-μm cell strainers, counted, and processed for downstream staining, enrichment, sorting, or functional assays.

### Human blood mononuclear cell preparation

Human blood mononuclear cells were isolated from cord blood and adult peripheral blood samples by Ficoll-Paque (#17144003, Cytiva) density-gradient centrifugation, washed, counted, and processed for downstream flow cytometric analysis.

### Flow cytometry and cell sorting

Single-cell suspensions were incubated with anti-CD16/CD32 Fc receptor block (clone 2.4G2; University of Michigan Hybridoma Core), before staining. In most experiments, anti-TCRγδ (#118118, Biolegend) was included in flow-cytometry panels, and CD8⁺ αβ T cells were defined as TCRβ⁺TCRγδ⁻. Intracellular staining was performed after fixation and permeabilization with the FOXP3/Transcription Factor Kit (#00-5523-00, Invitrogen). For lineage tracing analysis, cells were first fixed with IC Fixation Buffer (#50-112-9058, eBioscience) to preserve tdTomato fluorescence before permeabilization. Human peripheral blood and cord blood mononuclear cells were incubated with Human TruStain FcX (#422301, BioLegend) to block Fc receptors before staining. Antibodies used for mouse cell analysis, enrichment and/or sorting included anti-CD3 (17A2, #100214, Biolegend), anti-CD3ε (145-2C11, #100348, Biolegend), anti-TCRβ (H57-597, #109226, Biolegend), anti-Mouse TCR Vβ Screening Panel (#557004, BD Pharmingen), anti- TCRγδ (GL3, #118118, Biolegend), anti-CD45.1 (A20, #110723, Biolegend), anti-CD45.2 (104, #109841, Biolegend), anti-CD8α (53-6.7, #100761, Biolegend), anti-CD4 (GK1.5, #100428, Biolegend; RM4-5, #45004282, eBioscience), anti-CXCR3 (CXCR3-173, #126512, Biolegend), anti-CD44 (IM7, #103047, Biolegend), anti-CD49d (R1-2, #103617, Biolegend), anti-CD5 (53-7.3, #100625, Biolegend), anti-CD24 (M1/69, #101821, Biolegend), anti-CD69 (H1.2F3, #553237, BD Pharmingen), anti-CD103 (2E7, #121419, Biolegend), anti-NK1.1 (S17016D, #156510, Biolegend), anti-KLRG1 (2F1/KLRG1, #138416, Biolegend), anti-CD49b (DX5, #108922, Biolegend), anti-NKp46 (29A1.4, #137645, Biolegend), anti-SLAMF4 (m2B4(B6)458.1, #133503, Biolegend), anti-NKG2D (CX5, #130211, Biolegend), anti-Ly49A (YE1/48.10.6, #116805, Biolegend), anti-Ly49D (4E5, #555313, BD Pharmingen), anti-Ly49G2 (eBio4D11, #11578182, eBioscience), anti-Ly49H (3D10, #144713, Biolegend), anti-Ly49C/I (5E6, #562055, BD Pharmingen), anti-Asialo-GM1 (Poly21460, #146003, Biolegend), anti-CD11c (HL3, #558079, BD Pharmingen; N418, #12-0114-82, eBioscience), anti-CD11b (M1/70, #25011281, eBioscience), anti-IgM (RMM-1, #406503, Biolegend), anti-IgD (11-26c.2a, #405733, Biolegend), anti-CD124 (I015F8, #144805, Biolegend), anti-I-A/I-E (M5/114.15.2, #107607, Biolegend), anti-B220 (RA3-6B2, #103207, Biolegend), anti-CD25 (PC61.5, #25025181, eBioscience), anti-CD122 (TM-β1, #123221, Biolegend), anti-BCL-2 (BCL/10C4, #633504, Biolegend), anti-EOMES (W17001A, #157706, Biolegend), anti-T-bet (4B10, #644809, Biolegend), anti-PLZF (Mags.21F7, #53932080, eBioscience), anti-RORγt (B2D, #12698180, eBioscience), anti-Foxp3 (MF-14, #126403, Biolegend), anti-GATA3 (TWAJ, #12996642, eBioscience), anti-IFNγ (XMG1.2, #505825, Biolegend), anti-Granzyme B (QA16A02, #372203, Biolegend), anti-Ki67 (16A8, #652409, Biolegend), anti-rabbit IgG (Poly4064, #406419, Biolegend), and anti-Lin28b (EPR25252-20, #ab289978, Abcam). For Lin28b protein detection, cells were fixed and permeabilized before staining with anti-Lin28b followed by anti-rabbit IgG secondary antibody. Antibodies used for human flow cytometry included anti-TCRα/β (IP26, #306731, Biolegend), anti- CD8 (SK1, #341051, BD Pharmingen), anti-CD45RA (HI100, #304125, Biolegend), anti-NKG2A (REA110, #130-120-121, Miltenyi Biotec), anti-KIR3DL1/DL2 (REA970, #130-116-281, Miltenyi Biotec), anti-KIR2D (REA1042, #130-117-627, Miltenyi Biotec), anti-Asialo-GM1 (polyclonal, #53650780, eBioscience), and anti-EOMES (WD1928, #47487742, eBioscience). Antibodies were obtained from BioLegend, eBioscience, Abcam, BD Pharmingen, or Miltenyi Biotec. Data were acquired on BD FACSCanto (BD Biosciences) and BD LSRFortessa cytometers (BD Biosciences) and analyzed with FlowJo software (v10.10.0, FlowJo LLC). Cell sorting was performed on BD FACSAria III (BD Biosciences) and BD FACSDiscover S8 (BD Biosciences) instruments at the University of Michigan Flow Cytometry Core.

### Tetramer staining

Tetramer staining was performed using NP396-loaded MHC class I tetramer for LCMV-specific CD8⁺ αβ T cells, PBS-57-loaded CD1d tetramer for *i*NKT cells, and 5-OP-RU-loaded mouse MR1 tetramer for MAIT-cell detection. For MR1 tetramer staining, 6-FP-loaded mouse MR1 tetramer and *Mr1^⁻/⁻^* B6-MAIT^CAST^ samples were used as specificity controls. NP396-loaded MHC class I, PBS-57-loaded CD1d, and 5-OP-RU– and 6-FP–loaded MR1 tetramer reagents were provided by the NIH Tetramer Core Facility (NIH contract 75N93020D00005; RRID: SCR_026557; Atlanta, GA, USA).

### *Ex vivo* effector stimulation assays

Cultures were maintained at 37°C with 5% CO_2_. For TCR stimulation, splenocytes were cultured for 24 h using plate-bound anti-CD3 antibody (0.5-4 μg/ml; #100340, BioLegend), with or without soluble anti-CD28 antibody (1 μg/ml; #102116, BioLegend). GolgiStop (#512092KZ, BD Biosciences) was added during the final 4 h of culture. For cytokine stimulation, splenocytes were cultured for 24 h with IL-15 (50 ng/ml; #217-15, PeproTech), or IL-12 (5 ng/ml; #210-12, PeproTech) plus IL-18 (25 ng/ml; #B002-5, R&D Systems), with GolgiStop added during the final 4 h. For PMA/ionomycin stimulation, splenocytes were cultured for 4 h with PMA (50 ng/ml; # P8139, Sigma-Aldrich) and ionomycin (1 μg/ml; #I0634, Sigma-Aldrich) in the presence of GolgiStop. Human blood mononuclear cells were cultured for 24 h with human IL-15 (50 ng/ml; #200-15, PeproTech) with GolgiStop added during the final 4 h. Viable cells were identified using Fixable Viability Dye eFluor 450 (#65-0863-14, eBioscience) before flow cytometric analysis.

### Mixed lymphocyte reaction (MLR)

Negatively enriched splenic αβ T cells were used as responders. Cells were enriched by depletion of I-A/I-E⁺ (#107607, BioLegend), CD11c⁺ (#12-0114-82, eBioscience), CD11b⁺ (#101207, BioLegend), TCRγδ⁺ (#118107, BioLegend), B220⁺ (#103207, BioLegend), and NKp46⁺ (#137603, BioLegend) cells using the EasySep Mouse PE Positive Selection Kit II (#17666, STEMCELL Technologies), then labeled with CFSE (#30050, Biotium) according to the manufacturer’s instructions. CFSE-labeled responder cells were cultured with syngeneic or allogeneic splenocyte stimulators expressing a distinct congenic CD45 marker at a 2:1 responder-to-stimulator ratio. Cells cultured without stimulators served as negative controls, and cells stimulated with anti-TCRβ antibody (2 μg/ml; #109201, BioLegend) served as positive controls. Cultures were maintained at 37°C with 5% CO₂ for 4 days. Proliferation of responder CD8⁺ αβ T cells was assessed by CFSE dilution after gating on congenic-marker– defined responder TCRβ⁺TCRγδ⁻CD4⁻CD8α⁺ cells.

### Fetal and reaggregate thymus organ culture

For fetal thymic organ culture (FTOC), intact thymic lobes were isolated from embryonic day 15.5 embryos and cultured for 8 days at 37°C with 5% CO₂ on sponge-supported filter membranes composed of Surgifoam absorbable gelatin sponges (#ETH1972H, Medline) and Nuclepore track- etched membranes (#WHA10417301, Sigma-Aldrich). FTOC medium consisted of RPMI 1640 (#11875119, Gibco) supplemented with 10% FBS (#F4135, Sigma-Aldrich), 50 μM β- mercaptoethanol (#AB01340, AmericanBio), 10 mM HEPES (#15630106, Gibco), 1X nonessential amino acids (#11140050, Gibco), 1 mM sodium pyruvate (#11360070, Gibco), and 1X penicillin-streptomycin (#15140148, Gibco). Medium refreshed every 3-4 days. For reaggregate thymus organ culture (RTOC), CD45.2⁺ fetal thymic lobes were cultured with the presence of 1.35mM deoxyguanosine (dGuo; #D7145-25MG, Sigma-Aldrich) for 7 days to deplete endogenous thymocytes using the same culture scaffold as FTOC. Lobes were dissociated with 0.25% trypsin (#25200056, Gibco), and recovered thymic stromal cells were combined at a 1:1 ratio with sorted CD4⁻CD8⁻ DN thymocytes from adult CD45.1⁺ mice. After supernatant removal, cell pellets were gently resuspended to generate a concentrated cell slurry, and 0.5 −1 μl drops were placed onto the organ-culture scaffold. Reaggregates were cultured for 8 days using FTOC medium refreshed every 3–4 days.

### CyTOF staining, acquisition, and analysis

CyTOF antibody conjugation, staining, and data acquisition were performed as previously described^79^. Briefly, lanthanide metals (Standard BioTools, formerly Fluidigm) were conjugated to antibodies utilizing the Maxpar Antibody Labeling Kit (Standard BioTools). The following antibodies obtained from BioLegend, Invitrogen, Standard BioTools, Novus Biologicals, Abcam, or ThermoFisher were utilized for metal conjugation and staining: anti-CD45 (89Y-conjugated, 30- F11, #3089005B, Standard BioTools), anti-CD19 (112Cd-conjugated, 6D5, #Q10379, Invitrogen), anti-IFNγ (141Pr-conjugated, XMG1.2, #505812, Biolegend), anti-TNFα (142Nd-conjugated, MP6-XT22, #506302, Biolegend), anti-NKp46 (143Nd-conjugated, 29A1.4, #137625, Biolegend), anti-CD132 (144Nd-conjugated, TUGm2, #132306, Biolegend), anti-CD4 (145Nd-conjugated, RM4-5, #100520, Biolegend), anti-TCRβ (146Nd-conjugated, H57-597, #109235, Biolegend), anti-β2m (147Sm-conjugated, A16041A, #154502, Biolegend), anti-CD103 (148Nd-conjugated, 2E7, #121402, Biolegend), anti-CD8 (149Sm-conjugated, 53-6.7, #100716, Biolegend), anti- CD124 (150Nd-conjugated, I015F8, #144802, Biolegend), anti-CD49b (151Eu-conjugated, DX5, #108902, Biolegend), anti-Ly-6C (152Sm-conjugated, HK1.4, #NBP1-28046, Novus Biologicals), anti-CD122 (153Eu-conjugated, TM-B1, #123223, Biolegend), anti-CD11c (154Sm-conjugated, N418, #117302, Biolegend), anti-IA-IE (155Gd-conjugated, M5/114.15.2, #107602, Biolegend), anti-CD25 (156Gd-conjugated, 3C7, #101902, Biolegend), anti-CD27 (158Gd-conjugated, LG.3A10, #124202, Biolegend), anti-CD49a (159Tb-conjugated, HMα1, #142602, Biolegend), anti-IL-4 (160Gd-conjugated, 11B11, #504102, Biolegend), anti-IL-17a (161Dy-conjugated, TC11-18H10.1, #506935, Biolegend), anti-TCRγδ (162Dy-conjugated, GL3, #118101, Biolegend), anti-BCL-2 (163Dy-conjugated, BCL/10C4, #633502, Biolegend), anti-EOMES (164Dy-conjugated, Dan11mag, #14-4875-82, ThermoFisher), anti-CD49d (165Ho-conjugated, 9C10 (MFR4.B), #103708, Biolegend), anti-CXCR3 (166Er-conjugated, CXCR3-173, #126502, Biolegend), anti-CD127 (167Er-conjugated, A7R34, #135002, Biolegend), anti-T-bet (168Er- conjugated, 4B10, #644825, Biolegend), anti-Ki67 (169Tm-conjugated, 16A8, #652402, Biolegend), anti-CD62L (170Er-conjugated, MEL-14, #104402, Biolegend), anti-CD44 (171Yb- conjugated, IM7, #103014, Biolegend), anti-CD11b (172Yb-conjugated, M1/70, #3172012B, Standard BioTools), anti-CD69 (173Yb-conjugated, H1.2F3, #104502, Biolegend), anti-Sca-1 (174Yb-conjugated, D7, #108135, Biolegend), anti-KLRG1 (175Lu-conjugated, 2F1, #MAB69441-100, Novus Biologicals), anti-Granzyme B (176Yb-conjugated, GB11, #ab10912, Abcam), anti-CD3 (209Bi-conjugated, 145-2C11, #100314, Biolegend). Cells were stained with live-dead stain (Cisplatin-195Pt) (#201064, Standard BioTools), an antibody cocktail of cell surface markers, and an antibody cocktail of intracellular markers following fixation and permeabilization. Post-staining, cells were washed and stained with DNA Intercalator Iridium- 191/193 (#201192B, Standard BioTools) and stored at 4°C until acquisition on a CyTOF Helios system (Standard BioTools). EQ Four Element Calibration Beads (Standard BioTools) were added to the samples at the concentration recommended by the manufacturer prior to CyTOF acquisition. Data acquisition on the CyTOF system was conducted at a rate of approximately 100– 300 events per second. Post-acquisition, the instrument software applied a signal correction algorithm based on the calibration bead signals to adjust for any temporal variations in detector sensitivity. Immune cell subsets were annotated according to the following gating strategy: B cells (CD19^+^CD3^-^), Group 1 ILCs (CD3^-^CD49b^+^NKp46^+^), Myeloid cells (CD3^-^CD19^-^ CD11b^+^CD11c^+^CD49b^-^NKp46^-^), γδ T cells (CD3^+^TCRγδ^+^TCRβ^-^), CD4 T cells (CD3^+^TCRγδ^-^ TCRβ^+^CD4^+^CD8^-^), CD8 T cells (CD3^+^TCRγδ^-^TCRβ^+^CD4^-^CD8^+^).

### Single-cell RNA sequencing

Single cell suspensions were subjected to final cell counting on the Countess II Automated Cell Counter (ThermoFisher) and diluted to a concentration of 700 −1000 cells/ul. For FACS-sorted splenic αβ T cells (CD3⁺TCRγδ⁻TCRβ⁺) cells from WT and *β2m^⁻/⁻^MHCII^⁻/⁻^* mice, single cell 5’ gene expression and TCR libraries were generated using the 10x Genomics ChromiumX and following the manufacturer’s protocol for Chromium NextGEM Single Cell 5’ v2 Gene Expression reagents (10x Genomics). For FACS-sorted splenic WT IL-4-induced CXCR3^+^CD8^+^ αβ T cells (CD3^+^TCRβ^+^ TCRγδ^-^CD4^-^CD8^+^CXCR3^+^CD44^+^CD124^+^), splenic WT CXCR3⁻CD8⁺ αβ T cells (CD3^+^TCRγδ^-^ TCRβ^+^CD4^-^CD8^+^CXCR3^-^), WT CXCR3⁺CD8⁺ αβ T cells (CD3^+^TCRγδ^-^TCRβ^+^CD4^-^CD8^+^CXCR3^+^), and *β2m^⁻/⁻^MHCII^⁻/⁻^* CXCR3⁺CD8⁺ αβ T cells (CD3^+^TCRγδ^-^TCRβ^+^CD4^-^CD8^+^CXCR3^+^), 3’ single cell libraries were generated using the 10X Genomics Chromium Controller and following the manufacturer’s protocol for 3’ V3.1 chemistry with NextGEM Chip G reagents (10X Genomics). Final library quality was assessed using the Tapestation 4200 (Agilent) and libraries were quantified by Kapa qPCR (Roche). Pooled libraries were subjected to 150 bp paired-end sequencing according to the manufacturer’s protocol (Illumina NovaSeq 6000). Bcl2fastq2 Conversion Software v2.20 (Illumina) was used to generate de-multiplexed Fastq files and the CellRanger Pipeline (10X Genomics) was used to align reads and generate count matrices.

### Single-cell RNA sequencing data analysis

FACS-sorted splenic αβ T cells (CD3⁺TCRγδ⁻TCRβ⁺) cells from WT and *β2m^⁻/⁻^MHCII^⁻/⁻^* mice were processed for scRNA-seq and paired single-cell immune profiling of TCR repertoires. Gene- expression matrices and V(D)J clonotypes (TCR clonotypes) were generated by de-multiplexing, aligning, barcode processing, and UMI counting using Cell Ranger v6.1.2^80^. The R package Seurat (v4.2.0)^81^ was used to cluster the cells in the concatenated expression matrix. Cells expressing fewer than 100 genes, containing more than 2 x 10^4^ transcripts, or exhibiting >5% mitochondrial transcript content were excluded as low-quality cells. Gene expression values were normalized using NormalizeData with default parameters. The top 2000 variable genes were selected using FindVariableFeatures default, and counts were scaled and centered using ScaleData. Principal component analysis (PCA) was performed on these variable genes. The RunHarmony function from the R Harmony (V1.0)^82^ package was applied on the top 30 PCs to correct potential batch effects across individual samples. Uniform Manifold Approximation and Projection (UMAP) dimensional reduction was performed using RunUMAP with the top 20 Harmony dimensions. Cell clusters were identified using FindNeighbors and FindClusters, with the clustering resolution set to 0.5. Cluster marker genes were identified using FindAllMarkers, and differentially expressed genes between selected clusters were identified using FindMarkers. For independent scRNA-seq analysis of CXCR3-stratified CD8^+^ αβ T-cell subsets, FACS-sorted splenic WT CXCR3⁻CD8⁺ αβ T cells, WT CXCR3⁺ CD8⁺ αβ T cells, and *β2m^⁻/⁻^MHCII^⁻/⁻^* CXCR3⁺CD8⁺ αβ T cells were processed for scRNA-seq. Gene-expression matrices were generated using Cell Ranger v6.0.1. Seurat (v5.3.0)^83^ was used to load, merge, and cluster cells in the concatenated expression matrix. Cells expressing fewer than 200 genes or exhibiting >5% mitochondrial transcript content were excluded as low-quality cells. Gene expression values were normalized using NormalizeData with default parameters. Variable genes were selected using FindVariableFeatures with default parameters, and counts were scaled and centered using ScaleData. PCA was performed on variable genes. UMAP dimensional reduction was performed using RunUMAP. Cell clusters were identified using FindNeighbors and FindClusters, with the resolution set to 0.2. Cluster marker genes were identified using FindAllMarkers, and differentially expressed genes between selected clusters were found using FindMarkers. Data visualizations were performed in R version 4.4.2 (The R Foundation) and generated using variations of Seurat (v5.3.0)^83^ plotting functions, together with dplyr (v1.1.4)^84^, pheatmap (v1.0.13)^85^, ggplot2 (v3.5.2)^86^, ggrepel (v0.9.6)^87^, tidyr (v1.3.1)^88^, ggpubr (v0.6.0)^89^, multcompView (v0.1.10)^90^, RColorBrewer (v1.1.3)^91^, org.Mm.eg.db (v3.20.0)^92^, AnnotationDbi (v1.68.0)^93^, and enrichplot (v1.26.6)^94^. Gene set enrichment analysis (GSEA) was performed using clusterProfiler (v4.14.6)^95^ with Gene Ontology (GO) gene sets. A timestamp-derived neonatal score was calculated using the AddModuleScore function in Seurat (v5.3.0)^83^ based on genes significantly upregulated in early life-derived CD8⁺ αβ T cells timestamped at postnatal day 1 compared with adult life-derived CD8⁺ αβ T cells timestamped at postnatal day 28 from the publicly available dataset GSE97795^56^. The neonatal–adult developmental signature contained 462 genes, and the neonatal score was calculated from the 330 genes upregulated in day 1-timestamped CD8⁺ αβ T cells within this signature.

### Sequencing- and flow cytometry–based TCRβ repertoire profiling

For the splenic αβ T-cell scRNA-seq dataset, V(D)J gene-segment usage and CDR3 positional entropy were analyzed using scRepertoire (v2.2.1)^96^. TCR clonal diversity was quantified using the Shannon diversity index^97^, calculated as: *Index* = − ∑(*p_i_* × ln(*p_i_*)) where *p_i_* represents the proportional abundance of each TCR clonotype. For bulk TCRβ repertoire sequencing, sorted WT CXCR3⁻CD8⁺ αβ T cells, WT CXCR3⁺CD8⁺ αβ T cells, and *β2m^⁻/⁻^MHCII^⁻/⁻^* CXCR3⁺CD8⁺ αβ T cells were submitted to Adaptive Biotechnologies (Seattle, USA) for ImmunoSEQ analysis. V- and J-gene segment usage was visualized using the online immunoSEQ Analyzer platform. For flow cytometry–based TCR Vβ-chain usage analysis, splenic CD8⁺ αβ T cells from WT and *β2m^⁻/⁻^MHCII^⁻/⁻^* mice were stained with the Anti-Mouse TCR Vβ Screening Panel (#557004, BD Pharmingen) and analyzed after gating on CD3⁺TCRγδ⁻TCRβ⁺CD4⁻CD8α⁺ cells.

### Integrated scRNA-seq analysis

The integrative analysis comprises scRNA-seq data from FACS-sorted splenic CXCR3^-^CD8^+^ αβ T cells from WT mice, FACS-sorted splenic CXCR3^+^CD8^+^ αβ T cells from WT mice, FACS-sorted splenic CXCR3^+^CD8^+^ αβ T cells from *β2m^-/-^MHCII^-/-^*mice, and FACS-sorted splenic WT IL-4- induced CXCR3^+^CD8^+^ αβ T cell from IL-4C-treated WT mice. Quality control measures were applied to remove low-quality cells, excluding cells with fewer than 200 genes, more than 5,000 genes, or mitochondrial gene transcripts exceeding 25%. Ambient RNA was removed using the R package SoupX (v1.6.2)^98^ and doublets were removed using R package scDblFinder (v1.12.0)^99^. The data was normalized using the NormalizeData function and the “LogNormalize” method. The data was scaled using the ScaleData function and the top 2,000 most variable genes were identified using the FindVariableFeatures function. Dimensionality reduction was performed using RunPCA on the variable genes to derive principal components (PCs) and Uniform Manifold Approximation and Projection (UMAP) was applied to the top 50 PCs. Batch effect correction was conducted using R package Harmony (v1.0)^82^, with donor as the batch variable, and were used to generate a Uniform Manifold Approximation Projection (UMAP) representation of the data. Following batch correction, cells were clustered using the FindNeighbors function with shared nearest neighbor (SNN) modularity optimization-based clustering. Then the FindClusters function was performed for modularity optimization-based clustering to obtain biologically meaningful clusters.

### Public RNA-seq and scRNA-seq data processing and analysis

Public bulk RNA-seq and scRNA-seq datasets were obtained from the Gene Expression Omnibus (GEO) under accession numbers GSE112304^42^, GSE97795^56^, and GSE192480^28^. For bulk RNA- seq data, genes with fewer than 10 total counts across all samples were excluded. Differential expression analysis was performed using the DESeq2 (v1.46.0)^100^ package, applying a negative binomial generalized linear model. Genes were considered significantly differentially expressed if they exhibited an absolute log2 fold change greater than 0.6 and a Benjamini-Hochberg adjusted p-value (Padj) less than 0.05. For scRNA-seq analysis, datasets were processed using Seurat (v5.3.0)^83^. Samples were loaded, merged, and analyzed from the concatenated expression matrix. Cells with less than 100 genes or exhibiting >10% of mitochondrial transcript content were excluded as low-quality cells. Gene expression values were normalized using NormalizeData with default parameters. Variable genes were selected using FindVariableFeatures with default parameters, and counts were scaled and centered using ScaleData. PCA was performed on variable genes. UMAP dimensional reduction was performed using the RunUMAP. Cell clusters were identified using FindNeighbors and FindClusters, with the resolution set to 0.15. Cluster marker genes were found using FindAllMarkers, and differentially expressed genes between selected clusters were identified using FindMarkers. Data visualization was performed in R version 4.4.2 (The R Foundation), utilizing DESeq2 (v1.46.0)^100^ or Seurat (v5.3.0)^83^ plotting functions, together with dplyr (v1.1.4)^84^, pheatmap (v1.0.13)^85^, ggplot2 (v3.5.2)^86^, ggrepel (v0.9.6)^87^, and ggbreak (v0.1.4)^101^, and org.Mm.eg.db (v3.20.0)^92^ packages.

### Statistical analysis

All statistical analyses were performed using GraphPad Prism V10 (GraphPad Software). Differences between two groups were assessed using an unpaired two-tailed t-test. For comparisons involving more than two groups or multiple variables, one-way or two-way ANOVA was applied as appropriate. ANCOVA was used to assess statistically significant differences of simple linear regressions. A *p* value < 0.05 was considered statistically significant. Statistical significance is indicated as follows: \*\*\*\**p* < 0.0001, *** *p* < 0.001, ** *p* < 0.01, * *p* < 0.05; ns, not significant.

## KEY RESOURCE TABLE

| **REAGENT or RESOURCE** | SOURCE | IDENTIFIER |
|---|---|---|
| **Antibodies (anti-mouse)** |  |  |
| Rat monoclonal anti-CD3 | BioLegend | Cat#100214; RRID: AB_493645 |
| Armenian Hamster monoclonal anti-CD3ε | BioLegend | Cat#100348; RRID: AB_2564029 |
| Armenian Hamster monoclonal anti-TCRβ | BioLegend | Cat#109226; RRID: AB_1027649 |
| Armenian Hamster monoclonal anti-TCRγδ | BioLegend | Cat#118118; RRID: AB_10612756 |
| Mouse monoclonal anti-CD45.1 | BioLegend | Cat#110723; RRID: AB_493732 |
| Mouse monoclonal anti-CD45.2 | BioLegend | Cat#109807; RRID: AB_313444 |
| Rat monoclonal anti-CD8α | BioLegend | Cat#100761; RRID: AB_2564026 |
| Rat monoclonal anti-CD4 | BioLegend | Cat#100429; RRID: AB_493698 |
| Armenian Hamster monoclonal anti-CXCR3 | BioLegend | Cat#126512; RRID: AB_1088993 |
| Rat monoclonal anti-CD44 | BioLegend | Cat#103047; RRID: AB_2562451 |
| Rat monoclonal anti-CD49d | BioLegend | Cat#103617; RRID: AB_2563699 |
| Rat monoclonal anti-CD5 | BioLegend | Cat#100605; RRID: AB_312734 |
| Rat monoclonal anti-CD24 | BioLegend | Cat#101821; RRID: AB_756047 |
| Armenian Hamster monoclonal anti-CD69 | BD Pharmingen | Cat#553237; RRID: AB_394726 |
| Armenian Hamster monoclonal anti-CD103 | BioLegend | Cat#121419; RRID: AB_10709438 |
| Mouse monoclonal anti-NK1.1 | BioLegend | Cat#156510; RRID: AB_2876527 |
| Syrian Hamster monoclonal anti-KLRG1 | BioLegend | Cat#138416; RRID: AB_2561736 |
| Rat monoclonal anti-CD49b | BioLegend | Cat#108922; RRID: AB_2561460 |
| Rat monoclonal anti-NKp46 | BioLegend | Cat#137645; RRID: AB_2876479 |
| Mouse monoclonal anti-SLAMF4 | BioLegend | Cat#133503; RRID: AB_1595624 |
| Rat monoclonal anti-NKG2D | BioLegend | Cat#130211; RRID: AB_1236373 |
| Rat monoclonal anti-Ly49A | BioLegend | Cat#116805; RRID: AB_313756 |
| Rat monoclonal anti-Ly49D | BD Pharmingen | Cat#555313; RRID: AB_395724 |
| Rat monoclonal anti-Ly49G2 | eBioscience | Cat#11578182; RRID: AB_763604 |
| Mouse monoclonal anti-Ly49H | BioLegend | Cat#144707; RRID: AB_2561743 |
| Mouse monoclonal anti-Ly49C/I | BD Pharmingen | Cat#562055; RRID: AB_394750 |
| Armenian Hamster monoclonal anti-CD11c | BD Pharmingen | Cat#558079; RRID: AB_647251 |
| Rat monoclonal anti-CD11b | eBioscience | Cat#25011281; RRID: AB_469587 |
| Rat monoclonal anti-CD25 | eBioscience | Cat#25025181; RRID: AB_469607 |
| Rat monoclonal anti-CD122 | BioLegend | Cat#123221; RRID: AB_2750454 |
| Rat monoclonal anti-CD127 | BioLegend | Cat#135007; RRID: AB_1937231 |
| Rat monoclonal anti-CD215 | eBioscience | Cat#12714980; RRID: AB_10804884 |
| Mouse monoclonal anti-BCL-2 | BioLegend | Cat#633504; RRID: AB_2028394 |
| Rat monoclonal anti-EOMES | BioLegend | Cat#157706; RRID: AB_2888891 |
| Mouse monoclonal anti-T-bet | BioLegend | Cat#644809; RRID: AB_2028583 |
| Mouse monoclonal anti-PLZF | eBioscience | Cat#53932080; RRID: AB_2574445 |
| Rat monoclonal anti-RORγt | eBioscience | Cat#12698180; RRID: AB_10805392 |
| Rat monoclonal anti-Foxp3 | BioLegend | Cat#126403; RRID: AB_1089118 |
| Rat monoclonal anti-GATA3 | eBioscience | Cat#12996642; RRID: AB_1963600 |
| Rat monoclonal anti-IFNγ | BioLegend | Cat#505825; RRID: AB_1595591 |
| Mouse monoclonal anti-Granzyme B | BioLegend | Cat#372203; RRID: AB_2687027 |
| Rabbit polyclonal anti-Asialo-GM1 | BioLegend | Cat#146003; RRID: AB_2563195 |
| Rabbit monoclonal anti-Lin28b | Abcam | Cat#ab289978; RRID: N/A |
| Rat monoclonal anti-Ki67 | BioLegend | Cat#652409; RRID: AB_2562140 |
| Anti-Mouse TCR Vβ Screening Panel kit | BD Pharmingen | Cat#557004; RRID: AB_647180 |
| Rat monoclonal anti-IgM | BioLegend | Cat#406503; RRID: AB_315053 |
| Rat monoclonal anti-IgD | BioLegend | Cat#405733; RRID: AB_2563343 |
| Rat monoclonal anti-CD124 | BioLegend | Cat#144805; RRID: AB_2565598 |
| Rat monoclonal anti-I-A/I-E | BioLegend | Cat#107607; RRID: AB_313322 |
| Rat monoclonal anti-B220 | BioLegend | Cat#103207; RRID: AB_312992 |
| Fc receptor block (anti-CD16/CD32 clone 2.4G2) | This paper | N/A |
| **Antibodies (anti-rabbit)** |  |  |
| Donkey polyclonal anti-rabbit IgG | BioLegend | Cat#406419; RRID: AB_2563453 |
| **Antibodies (anti-human)** |  |  |
| Mouse monoclonal anti-TCRα/β | BioLegend | Cat#306731; RRID: AB_2650818 |
| Mouse monoclonal anti-CD8 | BD Pharmingen | Cat#341051; RRID: AB_400209 |
| Mouse monoclonal anti-CD45RA | BioLegend | Cat#304125; RRID: AB_10709440 |
| Human cell line monoclonal anti-NKG2A | Miltenyi Biotec | Cat#130-120-121; RRID: AB_2801749 |
| Human cell line monoclonal anti-KIR3DL1/DL2 | Miltenyi Biotec | Cat#130-116-281; RRID: AB_2727447 |
| Human cell line monoclonal anti-KIR2D | Miltenyi Biotec | Cat#130-117-627; RRID: AB_2733872 |
| Rabbit polyclonal anti-Asialo-GM1 | eBioscience | Cat#53650780; RRID: AB_10717257 |
| Mouse monoclonal anti-EOMES | eBioscience | Cat#47487742; RRID: AB_2573982 |
| **Tetramers (anti-mouse)** |  |  |
| PBS-57-loaded CD1d tetramer | NIH Tetramer Core Facility | N/A |
| 5-OP-RU-loaded mouse MR1 tetramer | NIH Tetramer Core Facility | N/A |
| 6-FP-loaded mouse MR1 tetramer | NIH Tetramer Core Facility | N/A |
| LCMV NP396 tetramer | NIH Tetramer Core Facility | N/A |
| **CyTOF antibodies (anti-mouse) and reagents** |  |  |
| Rat monoclonal anti-CD45 (89Y-conjugated) | Standard BioTools | Cat#3089005B; RRID: N/A |
| Mouse monoclonal anti-CD19 (112Cd-conjugated) | Invitrogen | Cat#Q10379; RRID: AB_10563403 |
| Rat monoclonal anti-IFNγ (141Pr-conjugated) | Biolegend | Cat#505812; RRID: AB_315406 |
| Rat monoclonal anti-TNFα (142Nd-conjugated) | Biolegend | Cat#506302; RRID: AB_315423 |
| Rat monoclonal anti-NKp46 (143Nd-conjugated) | Biolegend | Cat#137625; RRID: AB_2563744 |
| Rat monoclonal anti-CD132 (144Nd-conjugated) | Biolegend | Cat#132306; RRID: AB_2280163 |
| Rat monoclonal anti-CD4 (145Nd-conjugated) | Biolegend | Cat#100520; RRID: AB_312721 |
| Armenian Hamster monoclonal anti-TCRβ (146Nd-conjugated) | Biolegend | Cat#109235; RRID: AB_2562804 |
| Rat monoclonal anti-β2m (147Sm-conjugated) | Biolegend | Cat#154502; RRID: AB_2810419 |
| Armenian Hamster monoclonal anti-CD103 (148Nd-conjugated) | Biolegend | Cat#121402; RRID: AB_535945 |
| Rat monoclonal anti-CD8 (149Sm-conjugated) | Biolegend | Cat#100716; RRID: AB_312755 |
| Rat monoclonal anti-CD124 (150Nd-conjugated) | Biolegend | Cat#144802; RRID: AB_2561555 |
| Rat monoclonal anti-CD49b (151Eu-conjugated) | Biolegend | Cat#108902; RRID: AB_313409 |
| Rat monoclonal anti-Ly-6C (152Sm-conjugated) | Novus Biologicals | Cat# NBP1-28046; RRID: AB_1853354 |
| Rat monoclonal anti-CD122 (153Eu-conjugated) | Biolegend | Cat#123223; RRID: AB_2810372 |
| Armenian Hamster monoclonal anti-CD11c (154Sm-conjugated) | Biolegend | Cat#117302; RRID: AB_313771 |
| Rat monoclonal anti-IA-IE (155Gd-conjugated) | Biolegend | Cat#107602; RRID: AB_313317 |
| Rat monoclonal anti-CD25 (156Gd-conjugated) | Biolegend | Cat#101902; RRID: AB_312845 |
| Armenian Hamster monoclonal anti-CD27 (158Gd-conjugated) | Biolegend | Cat#124202; RRID: AB_1236456 |
| Armenian Hamster monoclonal anti-CD49a (159Tb-conjugated) | Biolegend | Cat#142602; RRID: AB_10915136 |
| Rat monoclonal anti-IL-4 (160Gd-conjugated) | Biolegend | Cat#504102; RRID: AB_315316 |
| Rat monoclonal anti-IL-17a (161Dy-conjugated) | Biolegend | Cat#506935; RRID: AB_2562850 |
| Armenian Hamster monoclonal anti-TCRγδ (162Dy-conjugated) | Biolegend | Cat#118101; RRID: AB_313826 |
| Mouse monoclonal anti-BCL-2 (163Dy-conjugated) | Biolegend | Cat#633502; RRID: AB_961014 |
| Rat monoclonal anti-EOMES (164Dy-conjugated) | ThermoFisher | Cat#14-4875-82; RRID: AB_11042577 |
| Rat monoclonal anti-CD49d (165Ho-conjugated) | Biolegend | Cat#103708; RRID: AB_2129607 |
| Armenian Hamster monoclonal anti-CXCR3 (166Er-conjugated) | Biolegend | Cat#126502; RRID: AB_1027635 |
| Rat monoclonal anti-CD127 (167Er-conjugated) | Biolegend | Cat#135002; RRID: AB_1937287 |
| Mouse monoclonal anti-T-bet (168Er-conjugated) | Biolegend | Cat#644825; RRID: AB_2563788 |
| Rat monoclonal anti-Ki67 (169Tm-conjugated) | Biolegend | Cat#652402; RRID: AB_11204254 |
| Rat monoclonal anti-CD62L (170Er-conjugated) | Biolegend | Cat#104402; RRID: AB_313089 |
| Rat monoclonal anti-CD44 (171Yb-conjugated) | Biolegend | Cat#103014; RRID: AB_312965 |
| Rat monoclonal anti-CD11b (172Yb-conjugated) | Standard BioTools | Cat#3172012B; RRID: AB_2661809 |
| Armenian Hamster monoclonal anti-CD69 (173Yb-conjugated) | Biolegend | Cat#104502; RRID: AB_313105 |
| Rat monoclonal anti-Sca-1 (174Yb-conjugated) | Biolegend | Cat#108135; RRID: AB_2563782 |
| Hamster monoclonal anti-KLRG1 (175Lu-conjugated) | Novus Biologicals | Cat#MAB69441-100; RRID: AB_3658681 |
| Mouse monoclonal anti-Granzyme B (176Yb-conjugated) | Abcam | Cat#ab10912; RRID: AB_297570 |
| Armenian Hamster monoclonal anti-CD3 (209Bi-conjugated) | Biolegend | Cat#100314; RRID: AB_312679 |
| DNA intercalator (191/193Ir-conjugated) | Standard BioTools | Cat#201192B; RRID: N/A |
| Live/Dead (Cisplatin) (195Pt-conjugated) | Standard BioTools | Cat#201064; RRID: N/A |
| **Bacterial and virus strains** |  |  |
| *Listeria monocytogenes* strain 10304S | Dr. Amale Laouar | N/A |
| LCMV strain Armstrong | Dr. Shannon Carty | N/A |
| **Chemicals, peptides, and recombinant proteins** |  |  |
| Phosphate Buffered Saline (PBS) | FISHER scientific | Cat#BP6651 |
| EasySep™ Mouse PE Positive Selection Kit II | STEMCELL technology | Cat#17666 |
| EasySep™ Mouse Streptavidin RapidSpheres™ Isolation Kit | STEMCELL technology | Cat#19860 |
| Peanut oil | Sigma-Aldrich | Cat#P2144 |
| Ficoll-Paque | Cytiva | Cat#17144003 |
| Murine IL-12 | Peprotech | Cat#210-12 |
| Murine IL-18 | R&D systems | Cat#B002-5 |
| Murine IL-15 | Peprotech | Cat#217-15 |
| Human IL-15 | Peprotech | Cat#200-15 |
| Murine IL-4 | Peprotech | Cat#214-14 |
| anti-mouse-IL-4 antibody | Bio X Cell | Cat#BE0045 |
| Golgi Stop | BD Biosciences | Cat#512092KZ |
| β-mercaptoethanol | Americanbio | Cat#AB01340 |
| Penicillin-Streptomycin (10,000 U/mL) | Gibco | Cat#15140148 |
| Fetal bovine serum (FBS) | Sigma-Aldrich | Cat# F4135 |
| Foxp3/Transcription Factor Staining Buffer Set | eBioscience | Cat# 00-5523-00 |
| IC fixation buffer | eBioscience | Cat# 50-112-9058 |
| Phorbol 12-myristate 13-acetate (PMA) | Sigma-Aldrich | Cat# P8139 |
| Ionomycin | Sigma-Aldrich | Cat#I0634 |
| Purified anti-mouse CD3ε | BioLegend | Cat#100340; RRID: AB_11149115 |
| Purified anti-mouse anti-CD28 | BioLegend | Cat#102116; RRID: AB_11147170 |
| Purified anti-mouse TCRβ | BioLegend | Cat#109201; RRID: AB_313424 |
| Agar | FISHER scientific | Cat#A360-500 |
| Brain Heart Infusion (BHI) Broth | Sigma-Aldrich | Cat#53286-100G |
| Poly(I:C) | Sigma-Aldrich | Cat#P1530-100MG |
| RPMI 1640 medium | Gibco | Cat#11875119 |
| HEPES (1M) | Gibco | Cat#15630106 |
| Sodium pyruvate (100mM) | Gibco | Cat#11360070 |
| MEM Non-Essential Amino Acids Solution (100X) | Gibco | Cat#11140050 |
| Deoxyguanosine (dGuo) | Sigma-Aldrich | Cat#D7145-25MG |
| Trypsin EDTA (0.25%) | Gibco | Cat#25200056 |
| CFSE Cell Proliferation Kits | Biotium | Cat#30050 |
| NAb Protein A/G Spin Kit | Thermo Fisher | Cat#89980 |
| Tamoxifen | Sigma-Aldrich | Cat#T5648 |
| Fixable Viability Dye eFluor™ 450 | eBioscience | Cat#65-0863-14 |
| Surgifoam Absorbable Gelatin Sponges | Medline | Cat#ETH1972H |
| Whatman® Nuclepore™ Track-Etched Membranes | Sigma-Aldrich | Cat#WHA10417301 |
| C1632 | Sigma-Aldrich | Cat#SML3673 |
| Human TruStain FcX | BioLegend | Cat#422301; RRID: AB_2818986 |
| DMSO | Sigma-Aldrich | Cat#D5879-500ML |
| 10% Formalin | VWR | Cat#77507-020 |
| **Deposited data** |  |  |
| CyTOF data: splenocytes | This paper | Zenodo repository accession number: 20722077 |
| Single-cell RNA-sequencing data: splenic αβ T cells | This paper | GEO: GSE335932 |
| Single-cell RNA-sequencing data: CXCR3-stratified CD8<sup>+</sup> αβ T cells | This paper | GEO: GSE335934 |
| Single-cell transcriptomic atlas of non-hematopoietic thymic compartments spanning fetal, neonatal, and adult stages | Gao et al.<sup>28</sup> | GEO: GSE192480 |
| Differentially expressed genes between 1 day and 28 days timestamped CD8 T cells | Smith et al.<sup>56</sup> | GEO: GSE97795 |
| Gene expression of naïve CD8 T cells, virtual memory T cells and true memory CD8 T cells | Quinn et al.<sup>42</sup> | GEO: GSE112304 |
| **Experimental models: Organisms/strains** |  |  |
| Mouse: C57BL/6J (B6) | The Jackson Laboratory | Strain #:000664; RRID: IMSR_JAX:000664 |
| Mouse: B6.SJL-*Ptprc<sup>a</sup> Pepc<sup>b</sup>*/BoyJ (CD45.1 B6) | The Jackson Laboratory | Strain #:002014; RRID: IMSR_JAX:002014 |
| Mouse: BALB/cJ (BALB/c) | The Jackson Laboratory | Strain #:000651; RRID: IMSR_JAX:000651 |
| Mouse: CByJ.SJL(B6)-*Ptprc<sup>a</sup>*/J (CD45.1 BALB/c) | The Jackson Laboratory | Strain #:006584; RRID: IMSR_JAX:006584 |
| Mouse: B6.129P2-*B2m<sup>tm1Unc</sup>*/DcrJ (*β2m<sup>−/−</sup>*) | The Jackson Laboratory | Strain #:002087; RRID: IMSR_JAX:002087 |
| Mouse: B6(129X1)-Tg(Cd4-cre/ERT2)11Gnri/J (CD4CreER<sup>T2</sup>) | The Jackson Laboratory | Strain #:022356; RRID: IMSR_JAX:022356 |
| Mouse: B6.Cg-*Gt(ROSA)26Sor<sup>tm9(CAG-tdTomato)Hze</sup>*/J (Ai9) | The Jackson Laboratory | Strain #:007909; RRID: IMSR_JAX:007909 |
| Mouse: B6.Cg-*Rag2<sup>tm1.1Cgn</sup>*/J (*Rag2<sup>-/-</sup>*) | The Jackson Laboratory | Strain #:008449; RRID: IMSR_JAX:008449 |
| Mouse: Nude *Foxn1<sup>−/−</sup>* | The Jackson Laboratory | Strain #:002019; RRID: IMSR_JAX:002019 |
| Mouse: C57BL/6NTac.Cg-*Rag2<sup>tm1Fwa</sup> Il2rg<sup>tm1Wjl</sup>* (*Rag2<sup>−/−</sup>Il2rγ<sup>−/−</sup>*) | Taconic Biosciences | Strain #:4111; RRID: IMSR_TAC:4111 |
| Mouse: B6.129-*H2-Ab1<sup>tm1Gru</sup> B2m<sup>tm1Jae</sup>* N17 (*β2m<sup>−/−</sup>MHCII<sup>−/−</sup>*) | Taconic Biosciences | Strain #:4080; RRID: IMSR_TAC:4080 |
| Mouse: CD45.1 *β2m<sup>−/−</sup>MHCII<sup>−/−</sup>* | Generated in the lab | N/A |
| Mouse: *Mr1<sup>+/+</sup>* B6-MAIT<sup>CAST</sup> | Dr. Olivier Lantz | Strain #: N/A<br>RRID: N/A |
| Mouse: *Mr1<sup>−/−</sup>* B6-MAIT<sup>CAST</sup> | Dr. Olivier Lantz | Strain #: N/A<br>RRID: N/A |
| **Software and algorithms** |  |  |
| Graphpad Prism 10 software | GraphPad Software, | RRID: SCR_002798<br>www.graphpad.com |
| FlowJo 10 | FlowJo LLC | RRID: SCR_008520 |
| R version 4.4.2 | The R Foundation | https://www.r-project.org/ |
| dplyr (v1.1.4) | Wickham et al.<sup>84</sup> | https://dplyr.tidyverse.org/ |
| DESeq2 (v1.46.0) | Love et al.<sup>100</sup> | https://bioconductor.org/packages/release/bioc/html/DESeq2.html |
| pheatmap (v1.0.13) | Kolde<sup>85</sup> | https://CRAN.R-project.org/package=pheatmap |
| ggplot2 (v3.5.2) | Wickham<sup>86</sup> | https://link.springer.com/book/10.1007/978-3-319-24277-4 |
| ggrepel (v0.9.6) | Slowikowski et al.<sup>87</sup> | https://CRAN.R-project.org/package=ggrepel |
| tidyr (v1.3.1) | Wickham et al.<sup>88</sup> | https://CRAN.R-project.org/package=tidyr |
| ggpubr (v0.6.0) | Kassambara<sup>89</sup> | https://CRAN.R-project.org/package=ggpubr |
| multcompView (v0.1.10) | Graves et al.<sup>90</sup> | https://CRAN.R-project.org/package=multcompView |
| RColorBrewer (v1.1.3) | Neuwirth<sup>91</sup> | https://CRAN.R-project.org/package=RColorBrewer |
| org.Mm.eg.db (v3.20.0) | Carlson<sup>92</sup> | https://bioconductor.org/packages/org.Mm.eg.db/ |
| AnnotationDbi (v1.68.0) | Pagès<sup>93</sup> | https://doi.org/10.18129/B9.bioc.AnnotationDbi |
| enrichplot (v1.26.6) | Yu et al.<sup>94</sup> | https://bioconductor.org/packages/enrichplot |
| clusterProfiler (v4.14.6) | Wu et al.<sup>95</sup> | https://doi.org/10.1038/s41596-024-01020-z |
| scRepertoire (v2.2.1) | Borcherding et al.<sup>96</sup> | https://doi.org/10.18129/B9.bioc.AnnotationDbi |
| Seurat (v4.2.0) | Hao et al.<sup>81</sup> | https://doi.org/10.1016/j.cell.2021.04.048 |
| Seurat (v5.3.0) | Hao et al.<sup>83</sup> | https://doi.org/10.1038/s41587-023-01767-y |
| Cell Ranger (v6.1.2) | Zheng et al.<sup>80</sup> | https://doi.org/10.1038/ncomms14049 |
| ggbreak (v0.1.4) | Xu et al.<sup>101</sup> | https://doi.org/10.3389/fgene.2021.774846 |
| Harmony (v1.0) | Korsunsky et al.<sup>82</sup> | https://cran.r-project.org/web/packages/harmony/index.html |
| SoupX (v1.6.2) | Young et al.<sup>98</sup> | https://doi.org/10.1093/gigascience/giaa151 |
| scDblFinder (v1.12.0) | Germain et al.<sup>99</sup> | https://doi.org/10.12688/f1000research.73600.2 |

## RESOURCE AVAILABILITY

### Lead contact

Further information and requests for resources and reagents should be directed to and will be fulfilled by the lead contact, Yasmina Laouar.

### Materials availability

This study did not generate new unique reagents.

### Data and code availability

- Data: Single-cell RNA sequencing data generated in this study have been deposited in the Gene Expression Omnibus under accession number GSE335935 (SuperSeries). Single-cell TCR immune profiling sequencing data generated in this study have been deposited in the Gene Expression Omnibus under accession number GSE335932. CyTOF data generated in this study have been deposited in the Zenodo repository under accession number 20722077.
- Code: This paper does not report original code.
- Any additional information required to reanalyze the data reported in this paper is available from the lead contact upon request.

